# Neural ensemble dynamics in dorsal motor cortex during speech in people with paralysis

**DOI:** 10.1101/505487

**Authors:** Sergey D. Stavisky, Francis R. Willett, Brian A Murphy, Paymon Rezaii, Donald T. Avansino, William D. Memberg, Jonathan P. Miller, Robert F. Kirsch, Leigh R Hochberg, A. Bolu Ajiboye, Krishna V. Shenoy, Jaimie M. Henderson

## Abstract

Speaking is a sensorimotor behavior whose neural basis is difficult to study at the resolution of single neurons due to the scarcity of human intracortical measurements and the lack of animal models. We recorded from electrode arrays in the ‘hand knob’ area of motor cortex in people with tetraplegia. Neurons in this area, which have not previously been implicated in speech, modulated during speaking and during non-speaking movement of the tongue, lips, and jaw. This challenges whether the conventional model of a ‘motor homunculus’ division by major body regions extends to the single-neuron scale. Spoken words and syllables could be decoded from single trials, demonstrating the potential utility of intracortical recordings for brain-computer interfaces (BCIs) to restore speech. Two neural population dynamics features previously reported for arm movements were also present during speaking: a large initial condition-invariant signal, followed by rotatory dynamics. This suggests that common neural dynamical motifs may underlie movement of arm and speech articulators.

## INTRODUCTION

Speaking requires coordinating numerous articulator muscles with exquisite timing and precision. Understanding how the sensorimotor system accomplishes this behavioral feat requires studying its neural underpinnings, which are critical for identifying (Tankus and Fried, 2018) and treating the causes of speech disorders and for building BCIs to restore lost speech (Guenther et al., 2009; Herff and Schultz, 2016). Speaking is also a uniquely human behavior, which presents a high barrier to electrophysiological investigations. Previous direct neural recordings during speaking have come from electrocorticography (ECoG) (Bouchard and Chang, 2014; Cheung et al., 2016; Mugler et al., 2014) or single-unit (SUA) recordings from penetrating electrodes during the course of clinical treatment for epilepsy (Chan et al., 2014; Creutzfeldt et al., 1989; Tankus et al., 2012) or deep brain stimulation for Parkinson’s disease (Lipski et al., 2018; Tankus and Fried, 2018). Such studies have begun to characterize motor cortical population dynamics underlying speech (Bouchard et al., 2013; Chartier et al., 2018; Pei et al., 2011), but not at the finer spatiotemporal scale uniquely afforded by high-density intracortical recordings, such as those available in animal models of reaching (Churchland et al., 2012; Kaufman et al., 2016; Miri et al., 2017).

We studied speech production at this resolution by recording from multielectrode arrays previously placed in human motor cortex as part of the BrainGate2 BCI clinical trial (Hochberg et al., 2006). This research context dictated two important elements of the present study’s design. First, both participants had tetraplegia due to spinal-cord injury but were able to speak; this enabled observing motor cortical spiking activity during overt speaking, in contrast to earlier studies of attempted speech by participants unable to speak (Brumberg et al., 2011; Guenther et al., 2009). However, these participants’ long-term paralysis means that their neurophysiology may differ from that of people who are able-bodied; we will discuss the need for interpretation caution in the Discussion.

Second, the electrode arrays were in dorsal ‘hand knob’ area of motor cortex which we previously found to strongly modulate to these participants’ attempted movement of their arm and hand (Ajiboye et al., 2017; Brandman et al., 2018; Pandarinath et al., 2017). Speech-related activity has not previously been reported in this cortical area, but there are several hints in the literature that dorsal motor cortex may have speech-related activity. Although imaging experiments consistently identify ventral cortical activation during speaking tasks, a meta-analysis of such studies (Guenther, 2016) indicates that responses are occasionally seen (though not, to our knowledge, explicitly called out) in dorsal motor cortex. Additionally, behavioral (Gentilucci and Campione, 2011; Vainio et al., 2013) and transcranial magnetic stimulation studies (Devlin and Watkins, 2007; Meister et al., 2003) have reported interactions (and interference) between motor control of the hand and mouth. This close linkage between hand and speech networks has been hypothesized to be due to a need for hand-mouth coordination and an evolutionary relationship between manual and articulatory gestures (Gentilucci et al., 2012; Rizzolatti and Arbib, 1998). Here, we explicitly set out to test whether neuronal firing rates in this dorsal motor cortical area modulated when participants produced speech and orofacial movements.

## RESULTS

### Speech-related activity in dorsal motor cortex

We recorded neural activity during speaking from participants ‘T5’ and ‘T8’, who previously had two arrays each consisting of 96 electrodes placed in the ‘hand knob’ area of motor cortex (**Figure 1A**). The participants performed a task in which on each trial they heard one of ten different syllables or one of ten short words, and then spoke the prompted sound after hearing a go cue (**Figure S1**). We analyzed both sortable SUA that could be attributed to an individual neuron’s action potentials, and ‘threshold-crossing’ spikes (TCs) that might come from one or several neurons (**Figure S2**). Firing rates showed robust changes during speaking of syllables (**Figures 1, S2**, **Supplemental Video 1**) and words (**Figure S7**). The neural population showed little modulation in the time epoch immediately after the audio prompt, prior to the go cue (**Figure S2C**). Since the response to the audio prompt was small, and since we are unable in this study to disambiguate between whether it reflects perception, movement preparation, or small overt movements preceding vocalization, we did not further examine this activity. Rather, here we focus on the neural activity leading up to and during speech production.

**Figure 1.**
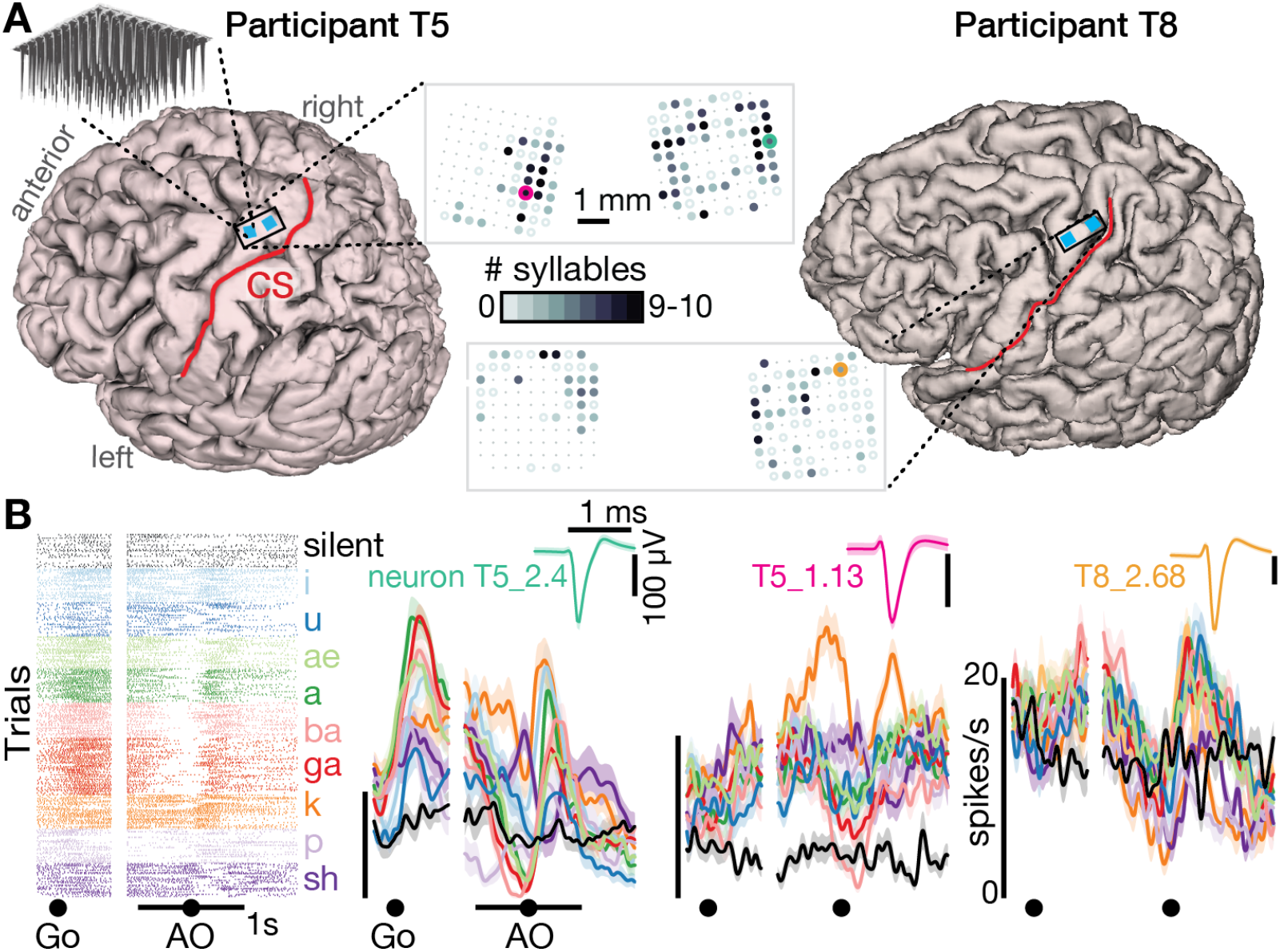
Speech-related neuronal spiking activity in dorsal motor cortex. (A) Participants’ MRI-derived brain anatomy. Blue squares mark the locations of the two chronic 96-electrode arrays. Insets show electrode locations, with shading indicating the number of different syllables for which that electrode recorded significantly modulated firing rates (darker shading = more syllables). Non-functioning electrodes are shown as smaller dots. CS is central sulcus. See also **Figure S2** for additional TCs firing rate examples, and **Figure S3** for individual syllables’ electrode response maps. **b.** Raster plot showing spike times of an example neuron across multiple trials of T5 speaking nine different syllables, or silence. Data are aligned to both the go cue and acoustic onset (AO). Trial-averaged firing rates for this neuron and two others are shown to the right (mean ± s.e.). Insets show these neurons’ action potential waveforms (mean ± s.d.). The electrodes where these neurons were recorded are circled in the panel A insets using colors corresponding to these waveforms. See **Figure S1** for task details.

Significant modulation was found during speaking at least one syllable (p < 0.05 compared to during silence) in 73/104 T5 electrodes’ TCs (13/22 SUA) and 47/101 T8 electrodes (12/25 SUA). Active neurons were distributed throughout the area sampled by the arrays, and most modulated to speaking multiple syllables (**Figures 1A and Figure S3**), suggesting a broadly distributed coding scheme. This is consistent with previous single neuron recordings in the temporal lobe (Creutzfeldt et al., 1989; Tankus et al., 2012). Two observations lead us to believe that this neural activity is related to the motor cortical control of the speech articulators (Chartier et al., 2018; Mugler et al., 2018) rather than perception or language. First, modulation was much stronger when speaking compared to hearing the auditory prompts (between 3.3x and 12.3x larger population responses, depending on the dataset; **Figure S2**). Second, in both participants, 99 of 120 electrodes that responded to syllables (24 of 25 sorted neurons) also responded to at least one of seven non-speech orofacial movements (**Figures 2 and Figure S4**). We also observed weak but significant firing rate correlation with breathing (**Figure S5**). Modulation for speaking was much stronger than for breathing (∼4.7x), and modulation for attempted arm movements was stronger than for speaking and orofacial movements (∼2.7x, **Figure S6**).

**Figure 2.**
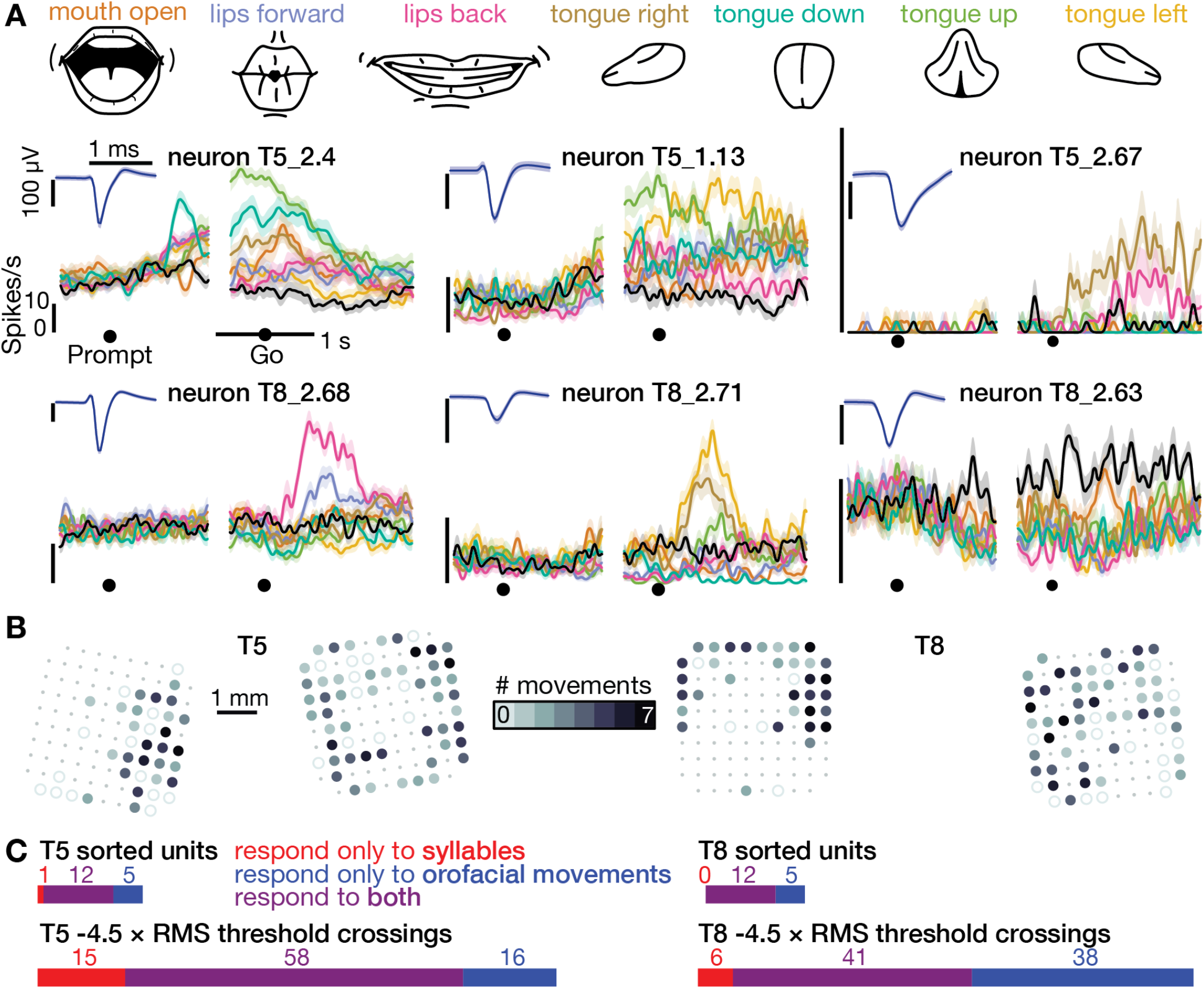
The same motor cortical population is also active during non-speaking orofacial movements. (A) Both participants performed an orofacial movement task during the same research session as their syllables speaking task. Examples of single neuron firing rates during seven different orofacial movements are plotted in colors corresponding to the movements in the illustrated legend above. The “stay still” condition is plotted in black. The same three example neurons from **Figure 1B** are included here. The other three neurons were chosen to illustrate a variety of observed response patterns. (B) Electrode array maps indicating the number of different orofacial movements for which a given electrode’s −4.5 × RMS threshold crossing rates differed significantly from the stay still condition. Data are presented similarly to the **Figure 1A** insets. Firing rates on most functioning electrodes modulated for multiple orofacial movements. (C) Breakdown of how many neurons’ (top) and electrodes’ TCs (bottom) exhibited firing rate modulation during speaking syllables only (red), non-speaking orofacial movements only (blue), or both behaviors (purple). A unit or electrode was deemed to modulate during a behavior if its firing rate differed significantly from silence/staying still for at least one syllable/movement.

### Speech can be decoded from intracortical activity on individual trials

We next performed a decoding analysis to quantify how much information about the spoken syllable or word was present in the time-varying neural activity. Multi-class support vector machines were used to predict the spoken sound (or silence) from single trial TCs and high-frequency LFP power (**Figure 3**). Cross-validated prediction accuracies for syllables were 84.6% for T5 (10 classes, mean chance accuracy was 0.1% across shuffle controls) and 54.7% for T8 (11 classes, chance was 8.6%). Word decoding accuracies were 83.5% for T5 (11 classes, chance was 9.1%) and 61.5% for T8 (11 classes, chance was 9.3%).

**Figure 3.**
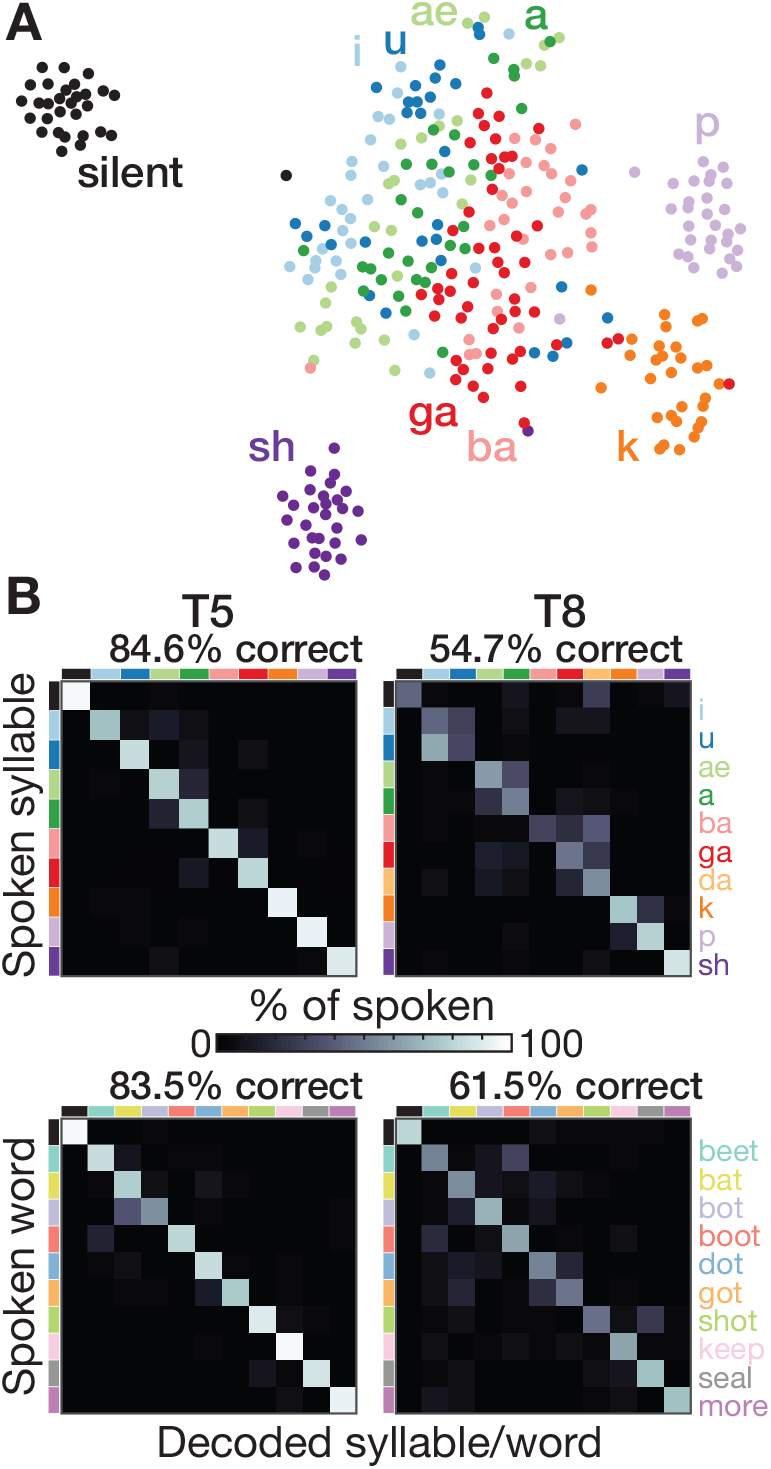
Speech can be decoded from intracortical activity. (A) To quantify the speech-related information in the neural population activity, we constructed a feature vector for each trial consisting of each electrode’s spike count and HLFP power in ten 100 ms bins centered on AO. For visualization, two-dimensional t-SNE projections of this feature vector are shown for all trials of the T5-syllables dataset. Each point corresponds to one trial. Even in this two-dimensional view of the underlying high-dimensional neural data, different syllables’ trials are discriminable and phonetically similar sounds’ clusters are closer together. (B) The high-dimensional neural feature vectors were classified using a multiclass SVM. Confusion matrices are shown for each participant’s leave-one-trial-out classification when speaking syllables (top row) and words (bottom row). Each matrix element shows the percentage of trials of the corresponding row’s sound that were classified as the sound of the corresponding column. Diagonal elements show correct classifications.

Decoding accuracies for all individual sounds were above chance (p < 0.01, shuffle test). Decoding mistakes (**Figure 3B**) and low-dimensional representations (**Figure 3A**) tended to follow phonetic similarities (e.g., *ba* and *ga*, *a* and *ae*). This observation is consistent with previous ECoG studies (Bouchard et al., 2013; Cheung et al., 2016; Mugler et al., 2014), although the larger neural differences we observed between unvoiced *k* and *p* and the beginning of their voiced counterparts at the start of *ga* and *ba* suggests strong laryngeal tuning (Dichter et al., 2018). These neural correlate similarities likely reflect similarities in the underlying articulator movements (Chartier et al., 2018; Lotte et al., 2015; Mugler et al., 2018).

### Neural population dynamics exhibit low-dimensional structure during speech

These multielectrode recordings enabled us to observe motor cortical dynamics during speech at their fundamental spatiotemporal scale: neuron spiking activity. Specifically, we examined whether two known key dynamical features of motor cortex firing rates during arm reaching were also present during speaking. Prior nonhuman primate (NHP) experiments showed that the neural state undergoes a rapid change during movement initiation which is dominated by a condition-invariant signal (CIS) (Kaufman et al., 2016). NHP (Churchland et al., 2012; Kaufman et al., 2016) and human (Pandarinath et al., 2015) studies found that subsequent peri-movement population activity is characterized by orderly rotatory dynamics. These observations, in concert with neural network modeling (Kaufman et al., 2016), have led to a model of motor control in which, prior to movement, inputs specifying the movement goal create attractor dynamics towards an advantageous initial condition (Shenoy et al., 2013). During movement initiation, a large transient input kicks the network into a different state from which activity evolves according to rotatory dynamics such that muscle activity is constructed from an oscillatory basis set (akin to composing an arbitrary signal from a Fourier basis set) (Churchland et al., 2012; Sussillo et al., 2015).

We tested whether motor cortical activity during speaking also exhibits these dynamics by applying the analytical methods of (Churchland et al., 2012; Kaufman et al., 2016). These analyses used two different dimensionality reduction techniques (Cunningham and Yu, 2014) to reveal latent low-dimensional structure in the trial-averaged firing rates for different conditions (here, speaking different words). Both methods sought to find a modest number of linear weightings of different electrodes’ firing rates (components) that capture a large fraction of the overall variance, akin to principal components analysis (PCA). However, unlike PCA, each method also looks for a specific form of neural population structure: jPCA (Churchland et al., 2012) seeks components with rotatory dynamics, whereas dPCA (Kaufman et al., 2016; Kobak et al., 2016) decomposes neural activity into CI and condition-dependent (CD) components. Importantly, these methods do not spuriously find the sought dynamical structure when it is not present in the data (Churchland et al., 2012; Elsayed and Cunningham, 2017; Kaufman et al., 2016; Kobak et al., 2016; Pandarinath et al., 2015).

We found that these population dynamics motifs were indeed also present during speaking. Similarly to (Kaufman et al., 2016), both participants’ neural activity featured a large CI component that rapidly increased after the go cue (**Figure 4A)**. This CIS_1_ was essentially identical regardless of which word was spoken (**Figure 4B**) and was largely orthogonal to the condition-dependent components.

**Figure 4.**
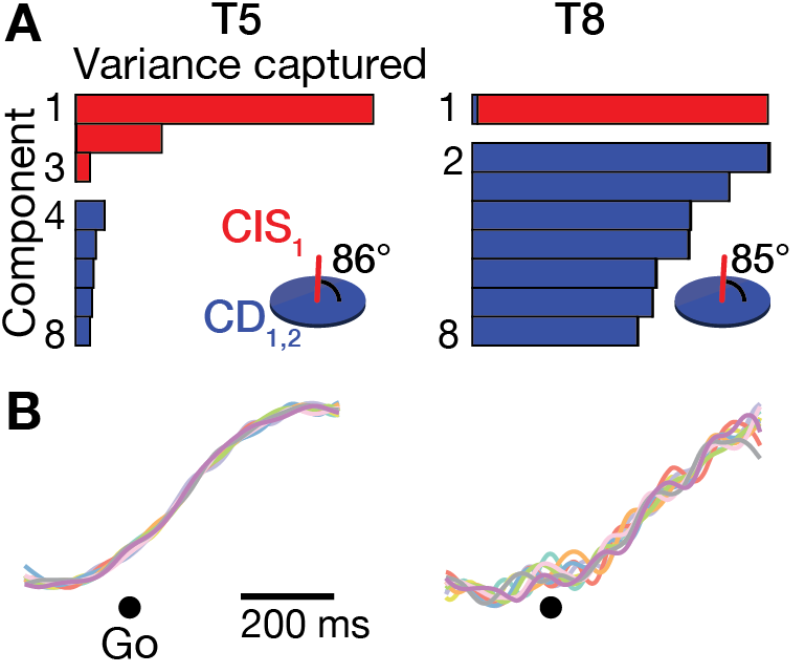
A condition-invariant signal during speech initiation. (A) A large component of neural population activity during speech initiation is a condition-invariant (CI) neural state change. Firing rates were decomposed into dPCA components like in (Kaufman et al., 2016). Each bar shows the relative variance captured by each dPCA component, which consists of both CI variance (red) and condition-dependent (CD) variance (blue). These 8 dPCs captured 65% (T5-words) and 32% (T8-words) of the overall neural variance. Insets show the subspace angle between the largest CI dimension (CIS_1_) and the two largest CD dimensions. Its angle to the subspace containing all the CD components was 82° for T5 and 73° for T8. (B) Neural population activity during speech initiation projected onto CIS_1_. Traces show the trial-averaged activity when speaking different words, denoted by the same colors as in **Figure 3B**.

We then looked for rotatory population dynamics around the time of acoustic onset. **Figure 5A** shows T5’s data projected into the top jPC plane. Similarly to (Churchland et al., 2012; Pandarinath et al., 2015), all conditions’ neural states rotated in the same direction, and rotatory dynamics could explain substantial variance in how population activity evolved moment-by-moment. Application of a recent population dynamics hypothesis testing method (Elsayed and Cunningham, 2017) revealed that this rotatory structure was significantly stronger than expected by chance in T5’s data (**Figure 5B**), but not T8’s (**Figure S8**). We attribute this difference to T8’s smaller measured neural responses during speech, which likely reflect his older arrays’ lower signal quality. Consistent with this, T8’s BCI computer cursor control performance was also substantially worse than T5’s (Pandarinath et al., 2017). Other factors that could also have contributed to T8’s reduced speech-related neural activity include his tendency to speak quietly and with less clear enunciation (consistent with (Jiang et al., 2016)), array placement differences, and differences in cortical maps between individuals (Farrell et al., 2007).

**Figure 5.**
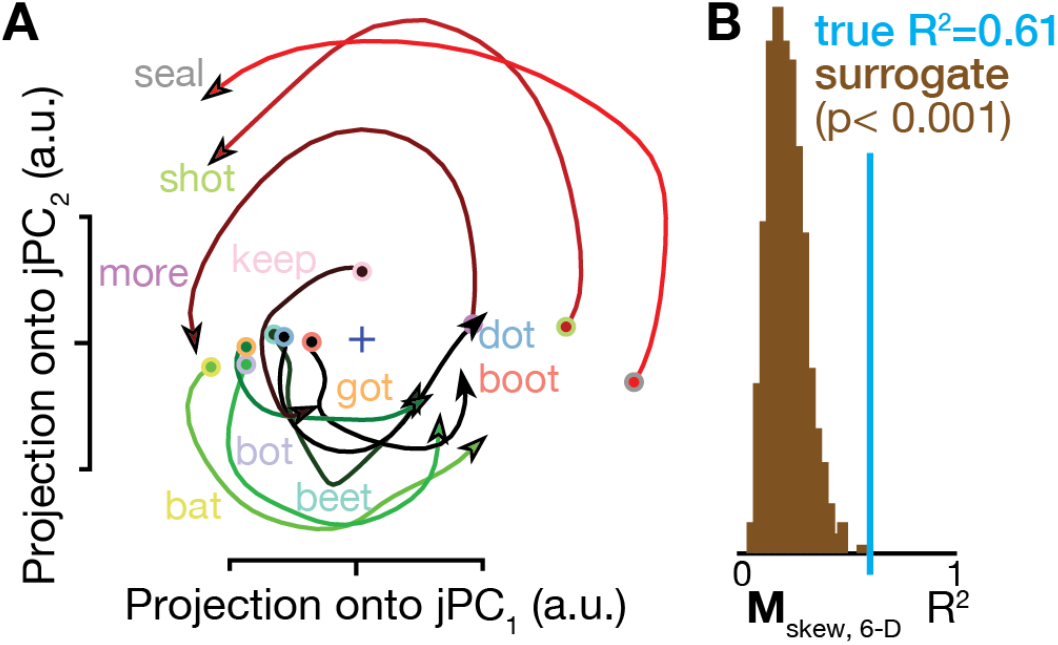
Rotatory neural population dynamics during speech. (A) The top 6 PCs of the trial-averaged firing rates from 150 ms before to 100 ms after acoustic onset in the T5-words dataset were projected onto the first jPCA plane like in (Churchland et al., 2012). This plane captures 38% of the top 6 PCs’ variance, and rotatory dynamics fit the moment-by-moment neural state change with R^2^ = 0.81 in this plane and 0.61 in the top 6 PCs. See also **Figure S8** for participant T8’s rotatory dynamics results. (B) Statistical significance testing of rotatory neural dynamics during speaking. The blue vertical line shows the goodness of fit of explaining the evolution in the top 6 PC’s neural state from moment to moment using a rotatory dynamical system. The brown histograms show the distributions of this same measurement for 1,000 neural population control surrogate datasets generated using the tensor maximum entropy method of (Elsayed and Cunningham, 2017). These shuffled datasets serve as null hypothesis distributions that have the same primary statistical structure (mean and covariance) as the original data across time, electrodes, and word conditions, but not the same higher-order statistical structure (e.g., low-dimensional rotatory dynamics).

## DISCUSSION

There are three main conclusions from these findings. First, they suggest that ‘hand knob’ motor cortex, an area not previously known to be active during speaking (Breshears et al., 2015; Dichter et al., 2018; Leuthardt et al., 2011; Lotte et al., 2015), may in fact play a role in speech production. Speech-related single-neuron modulation might have been missed by previous studies due to the coarser resolution of ECoG (Chan et al., 2014). If this finding holds true in the wider population, this would underscore that the familiar ‘motor homunculus’ (Penfield and Boldrey, 1937) is overly simplistic. While it is generally recognized that motor cortex does not follow a sequential point-to-point somatotopy (and indeed, Penfield and colleagues were aware of this and intended for their diagram to be a simplified overview), the patchy mosaicism amongst smaller parts in the current view of precentral gyrus organization still features a dorsal-to-ventral progression and separation of the major body regions (leg, arm, head) (Farrell et al., 2007; Schieber, 2001). The presence of neurons responding to face and tongue movements in the dorsal “arm/hand” area of motor cortex could indicate that sensorimotor maps for different body parts are even more widespread and overlapping than previous thought. Given our previous finding that activity from these same arrays encodes intended arm and hand movements (Pandarinath et al., 2017), these observations would also support the hypothesis that the systems for speech and manual gestures are interlocked (Gentilucci et al., 2012; Rizzolatti and Arbib, 1998; Vainio et al., 2013).

An important unanswered question, however, is to what extent these results were potentially influenced by cortical remapping due to tetraplegia. While we cannot rule this out, we believe that remapping of face representation to the hand knob area is unlikely. Despite these participants’ many years of paralysis, the sites we recorded from still strongly respond to attempted hand and arm movements (Ajiboye et al., 2017; Brandman et al., 2018; Pandarinath et al., 2017), and we verified in participant T5 that modulation during attempted arm movements was stronger than during speech production. These results are inconsistent with this area being “taken over” by functions related to orofacial movements. Furthermore, motor cortical remapping following arm amputation was recently shown to be smaller than previously thought (Wesselink et al., 2019), and in particular much smaller than what would be needed to move lip representations to hand cortex (Makin et al., 2015). Definitively resolving this ambiguity would require intracortical recording from this eloquent brain area in able-bodied people.

Second, the offline decoding results demonstrate the potential utility of using intracortical signals to restore speech to people with some forms of anarthria by transforming their intended speech into audible sounds or text. Decoding the neural correlates of attempted speech production (Brumberg et al., 2011) may be more desirable over approaches that decode covert internal speech (Leuthardt et al., 2011; Martin et al., 2016) or more abstract elements of language (Chan et al., 2011; Yang et al., 2017) because it leverages existing neural machinery that separates internal monologue and speech preparation from intentional speaking. The present results compare favorably to previously published decoding accuracies using ECoG (Mugler et al., 2014; Ramsey et al., 2018) despite our dorsal recording locations likely being suboptimal for decoding speech. Multi-electrode arrays placed in ventral motor cortex would be expected to yield even better decoding accuracies. Furthermore, recent order-of-magnitude advances in the number of recording sites on intracortical probes (Jun et al., 2017) point to a path that stretches far forward in terms of scaling the number of distinct sources of information (neurons) for speech BCIs. That said, the present results are only a first step in establishing the feasibility of speech BCIs using intracortical multielectrode arrays. Here we decoded amongst a limited set of discrete syllables and words in participants who are able to speak; future studies will be needed to assess how well intracortical signals can be used to discriminate between a wider set of phonemes (Brumberg et al., 2011; Mugler et al., 2014), in the absence of overt speech (Brumberg et al., 2011; Martin et al., 2016), and to synthesize continuous speech (Anumanchipalli et al., 2018).

Third, we showed that two motor cortical population dynamics motifs present during arm movements — a large condition-invariant change at movement initiation and rotatory dynamics during movement generation – were also significant features of speech activity. We speculate that these neural state rotations are well-suited for generating descending muscle commands driving the out-and-back articulator movements that form the kinematic building blocks of speech (Chartier et al., 2018; Mugler et al., 2018). Future research involving recording from the relevant muscles (Churchland et al., 2012) and causally stimulating the circuit (Dichter et al., 2018) is needed to test this hypothesis. The presence of these dynamics during both reaching and speaking could indicate a conserved computational mechanism that is ubiquitously deployed across behaviors to shift the circuit dynamics from withholding movement to generating the appropriate muscle commands from an oscillatory basis set.

## ACKNOWLEDGEMENTS

We thank participants T5, T8, and their caregivers for their dedicated contributions to this research, N. Lam for administrative support, and Dr. Sydney Cash and Dr. Laura Ball for helpful discussions.

## Author contributions

S.D.S. designed the study and wrote the manuscript. S.D.S., B.A.M., P.R., F.R.W., and D.T.A collected the data. S.D.S. and F.R.W. analyzed the data. L.R.H. is the sponsor-investigator of the multi-site clinical trial. J.M.H. and J.P.M. planned and performed the array placement surgeries for T5 and T8, respectively, and were responsible for their ongoing clinical care. W.D.M. assisted in T8’s recruitment, surgery, and his day-to-day research activities. A.B.A and R.F.K. supervise and are responsible for the clinical site where T8 was enrolled. J.M.H. and K.V.S. guided the study and supervise and are responsible for the clinical site where T5 was enrolled. All authors reviewed and edited the manuscript.

## Materials and data

Data may be made available upon reasonable request to the senior authors (K.V.S. or J.M.H.). Please note that sharing of raw human neural data is restricted due to the potential sensitivity of this data in combination with the very small number of BrainGate2 trial participants. Code may be made available upon request to the corresponding author (S.D.S.).

## METHODS

### Participants

The two participants in this study were enrolled in the BrainGate2 Neural Interface System pilot clinical trial (ClinicalTrials.gov Identifier: NCT00912041). The overall purpose of the study is to obtain preliminary safety information and demonstrate proof of principle that an intracortical brain-computer interface can enable people with tetraplegia to communicate and control external devices. Permission for the study was granted by the U.S. Food and Drug Administration under an Investigational Device Exemption (Caution: Investigational device. Limited by federal law to investigational use). The study was also approved by the Institutional Review Boards of Stanford University Medical Center (protocol #20804), Brown University (#0809992560), University Hospitals of Cleveland Medical Center (#04-12-17), Partners HealthCare and Massachusetts General Hospital (#2011P001036), and the Providence VA Medical Center (#2011-009). Both participants gave informed consent to the study and publications resulting from the research, including consent to publish photographs and audiovisual recordings of them.

**Participant ‘T5’** (male, right-handed, 64 years old at the time of the study) was diagnosed with C4 AIS-C spinal cord injury ten years prior to these research sessions. He retained the ability to weakly flex his left elbow and fingers and some slight and inconsistent residual movement of both the upper and lower extremities. T5 was able to speak normally and converse naturally without hearing assistance, but had some trouble hearing from his left ear.

**Participant ‘T8’** (male, right-handed, 56 years old at the time of the study) was diagnosed with C4 AIS-A spinal cord injury eleven years prior to these sessions. He retained restricted and non-functional voluntary shoulder girdle motion on both sides, and non-functional voluntary finger extension on his left side. He had no sensation below the shoulder. T8 was able to speak normally and converse naturally with the assistance of hearing aids in both his ears.

### Prompted speaking tasks

Participants performed a **syllables task** consisting of discrete trials in which they spoke out loud one of ten different phonemes or consonant-vowel syllables in response to an auditory prompt. These prompts were *i* (as in “beet”); *ae* (as in “bat”); *a* (as in “bot”); *u* (as in “boot”); *ba; da; ga; sh* (as in the start of “shot”), and the unvoiced *k* and *p*. All pronunciations were American English. **Supplemental Video 1** provides a continuous audio recording of one set of each type of syllables task trial.

Participants sat comfortably in a chair facing a microphone in a quiet room. They were instructed to refrain from attempting movements or speaking during trials except when prompted to speak by a custom experiment control software written in MATLAB (The Mathworks, USA). During trials they were also asked to fixate on the same object in front of them. A trial began with two beeps to alert the participant that the trial was starting. After 0.4 seconds, a pre-recorded syllable prompt was played via computer speakers. After 0.8 seconds, two clicks served as the go cue that instructed the participant to speak back the prompted sound. The next trial started 2.2 seconds later. There was also an eleventh ‘silent’ condition which was identical to the spoken syllables trials, except that instead of playing a syllable prompt, the speakers played a nearly-silent audio file consisting of ambient background noise recorded in the same environment as the syllable prompts. The participants had been previously instructed not to say anything in response to this silent prompt.

The task was performed in blocks consisting of ten trial sets. Each set contained eleven trials: one trial of each syllable, plus silence, presented in a randomized order. After the task was explained to each participant, he was given time to practice a few sets of the task until he indicated that he was ready to begin data collection. At the end of each set we paused the task until the participant indicated that he was ready to continue. These inter-set pauses typically lasted less than ten seconds. Participants performed three consecutive blocks of the task during a research session, with longer pauses of several minutes between blocks during which we encouraged the participant to rest, adjust his posture for comfort, and take a drink of water.

Both the audio prompts played by the experiment control computer, and the participant’s voice, were recorded by the microphone (Shure SM-58). This audio signal was recorded via the analog input port of the electrophysiology data acquisition system and digitized at 30 ksps together with the raw neural data (see Neural Recording section). Each trial’s acoustic onset time (AO) was manually determined by visual and auditory inspection of the recorded audio data. During this review, we also excluded infrequent trials where the participant spoke at the wrong time or when the trial was interrupted (for example, if a caregiver entered the room). Isolated sounds can be difficult to discriminate, and our participants sometimes misheard a syllable prompt as a phonetically similar prompt. In particular, T5 misheard the majority of *da* as *ga* (or occasionally as *ba*). Both participants made a few other substitutions between similar syllables. In this study we were interested in the neural correlates of preparing and then generating speech, which should reflect the syllable that the participant perceived. We therefore labeled these misheard trials based on the spoken, rather than prompted, syllable for subsequent analyses. This left an insufficient number of T5 *da* trials for subsequent neural analyses; thus, there are eleven conditions shown in T8’s **Figure 1B** firing rate plots and **Figure 3B** confusion matrices, but only ten conditions for T5. The number of trials analyzed for each participant, after excluding trials and re-labeling misheard trials as described above, were: silent (30 trials for T5, 30 trials for T8); *i* (30, 28); *u* (30, 31); *ae* (28, 30); *a* (30, 30); *ba* (31, 29); *ga* (50, 34); *da* (0, 27); *k* (30, 27); p (30, 33); *sh* (30, 30). We refer to these datasets as ‘T5-syllables’ and ‘T8-syllables’.

Participants also performed a **words task** which was identical to the syllables task except that they repeated back one of ten short words, rather than syllables, in response to the auditory prompt. Each participant performed three blocks of ten repetitions of each word during one research session. We refer to these datasets as ‘T5-words’ and ‘T8-words’. Two consecutive trials were excluded from the T8-words dataset because of a large electrical noise artifact across almost all electrodes. The specific words, and the number of trials analyzed for each participant, were: “beet” (30 T5 trials, 29 T8 trials); “bat” (30, 29); “bot” (30, 28); “boot” (30, 30); “dot” (30, 29); “got” (29, 29); “shot” (29, 28); “keep” (30, 30); “seal” (30, 30); “more” (30, 30). As with the syllables task, there was also a silent condition (30 T5 trials, 30 T8 trials).

Silent condition trials were assigned a ‘faux AO’ so that neural data from comparable epochs of silent and spoken trials could be visualized and analyzed (for example, for generating trial-averaged, AO-aligned firing rates in **Figure 1** or for decoding silent trials’ neural activity in **Figure 3**). Specifically, each silent trial’s AO was set to equal the mean AO (relative to the go cue) for all the spoken syllables or words during the same block.

### Orofacial movement task

Participants also performed an orofacial movement task with a similar trial structure as the syllables and words tasks. Seven different movement conditions were instructed with auditory prompts: “mouth open”, “lips forward”, “lips back”, “tongue right”, “tongue down”, “tongue up”, and “tongue left”. An additional “stay still” condition was analogous to the silent condition of the syllables and words tasks. Prior to the first block of the orofacial task, a researcher explained the prompts to the participant, demonstrated the movements, and ran the participant through a few practice sets. Due to clinical trial protocols, we did not collect kinematic tracking data such as electromagnetic midsagittal articulography (Chartier et al., 2018) or ultrasound recordings. A video recording of the participants’ faces (without markers) did allow the researchers to confirm that the participants were making the instructed movement with acceptable timing precision. Given this limitation, we limited our use of these data to broadly testing for neural responses during orofacial movements, rather than quantifying precise moment-by-moment relationships between neural activity and kinematics.

Similar to the syllables and words task, a trial began with two ready beeps, after which the computer speaker played a movement prompt (e.g., “lips forward”). This was followed by the pair of go clicks; the participants were previously informed that they should begin moving after the second click. 1.3 seconds later, the experiment control system played the verbal command “return”, which instructed the participant to return to a neutral orofacial posture (e.g., close the mouth after “mouth open”, move the tongue left after “tongue right”). The trial ended 1.2 seconds later. The purpose of using a return cue was so that there was a known epoch after the movement go cue during which we knew that the participant was not yet returning. The return cue also provided the participant with dedicated time to return to a neutral orofacial position, so that all trials would start from roughly the same posture. For T8, the “return” instruction was immediately followed by a go click. However, we observed that T8 started the return movement upon hearing “return” rather than waiting for the go click. We therefore removed the return go click prior to T5’s research sessions, and instead instructed T5 to start the return movement when he heard “return”. In the present study we did not examine the return portion of the orofacial movement task.

Each participant’s orofacial movements and syllables datasets were collected on the same day during the same research session; three blocks of the orofacial movements task immediately followed three blocks of the syllables task. We will refer to these orofacial movements task datasets as ‘T5-movements’ and ‘T8-movements’. No trials were excluded from these datasets; thus, there were 30 trials of each condition for each participant.

### Breath measurement

T5’s breath-related abdomen movements were measured with a MLT1132 piezo respiratory belt transducer (model MLT1132, ADIntruments, USA). The stretch sensor was wrapped around his abdomen at the point where it maximally expanded during breathing. Analog voltage signals from the belt were input to the neural signal processor via one of its analog input channels. These data were digitized at 30 ksps along with the neural data. Our goal was to test whether there is breath-correlated neural activity during natural “background” breathing, when the participant was not consciously attending to their breath. We therefore recorded neural and breath proxy measurements while he performed a BCI computer cursor task as part of a different set of experiments, and during an interval where he was resting quietly after completing the BCI task. We refer to this as the ‘t5-breathing’ dataset.

### Movement comparisons task

The purpose of this task, which was performed on a separate day, was to compare the neural modulation when making orofacial movements and speaking, versus when attempting to make arm and hand movements. The task had a visually-instructed delay structure: during the instructed delay period, a red square appeared in the center of a computer monitor facing the participant. Text displayed above the square cued the upcoming movement, for example, “Prepare: Say Ba”, or “Prepare: Open Hand”. There was also a “Prepare: Do Nothing” instruction condition, which otherwise had the same trial structure as the instructed movements. After a random delay period of between 1400 and 1800 ms, a go cue appeared; this consisted of the center square turning from red to green, and the text changing to “Go”. During this epoch, T5 attempted to make the instructed movement as best as he could. This resulted in complete movements for all the orofacial and speaking movements and “shoulder shrug”, partial movements for some of the arm movements (e.g., “flex elbow in”), and no overt movement for the other arm movements (e.g., “close hand”, “thumb up”). We analyzed neural data from 200 ms to 600 ms after the go cue. We note that insofar as there was somatosensory and proprioceptive feedback only during the actualized movements, this would be expected to increase the observed neural modulation to orofacial movements and speaking, and decrease the modulation to attempted arm and hand movements. The go cue stayed on for 1500 ms. This was followed by a return period in which the text changed to “Return”; during this epoch, the participant was instructed to return his body to a neutral posture. 32 trials were collected for each movement type. We refer to this as the ‘t5-comparisons’ dataset.

### Neural recording

Both participants had two 96-electrode Utah arrays (1.5 mm electrode length, Blackrock Microsystems, USA) neurosurgically placed in dorsal ‘hand knob’ area of the left (motor dominant) hemisphere’s motor cortex. T5 and T8 had the arrays placed 14 and 34 months prior to the present study’s speaking tasks, respectively. The t5-breathing and t5-comparisons datasets were recorded 26 months after array placement. Arrays were placed in areas anticipated to have arm movement-related activity because two goals of the clinical trial are 1) testing the feasibility of intracortical BCI-based communication using point-and-click keyboards and 2) restoration of reach and grasp function via control of a robotic arm or functional electrical stimulation. We note that these implant sites are distinct from the closest known speech area, which is the dorsal laryngeal motor cortex (Bouchard et al., 2013; Dichter et al., 2018). In this study, we looked for neural correlates of speaking in dorsal motor cortex. To help contextualize the results, here we summarize the other intended behaviors associated with modulation of the neural activity recorded by these same arrays. Our previous studies have reported that T5 and T8 controlled BCI computer cursors by attempting movements of their arm and hand (Brandman et al., 2018; Pandarinath et al., 2017). T8 was also able to use intended arm movements to command movements of his own paralyzed arm via functional electrical stimulation (Ajiboye et al., 2017). We also recorded movement task outcome error signals from T5’s arrays; these signals indicated whether the participant succeeded or failed at acquiring a target using a BCI-controlled cursor (Even-Chen et al., 2018).

Neural signals were recorded from the arrays using the NeuroPort™ system (Blackrock Microsystems). Voltage was measured between each of the 96 electrodes’ uninsulated tips and that array’s reference wire. Wire bundles ran from each array to cranially-implanted connector pedestals. During research sessions, a ‘patient cable’ with a unity gain pre-amplifier was connected to each array’s corresponding pedestal and carried signals to an isolated unity gain front-end amplifier. These signals were analog filtered from 0.3 Hz to 7.5 kHz, digitized at 30 kHz (250 nV resolution), and sent to the neural signal processor via fiber-optic link. As mentioned earlier, amplified analog voltage data from the microphone were input to the neural signal processor and were digitized time-locked with the neural signals. All of these digitized data were sent over a local network to a connected PC where they were recorded to hard disk for subsequent analysis.

The naming scheme for neurons or electrodes in figures is <participant>_<array #>.<electrode #>. For example, “neuron T5_2.4” in **Figure 1** refers to a participant T5 neuron identified on the second array (which is the more medial of each participant’s two arrays) on electrode #4 (according to the manufacturer’s electrode numbering scheme).

### Neural signal processing

Neuronal action potentials (spikes) were detected as follows. We first applied a common average re-referencing to each electrode within an array by subtracting, at each time sample, the mean voltage across all electrodes on that array. These voltage signals were then filtered with a 250 Hz asymmetric FIR high pass filter designed to extract spike activity from this type of array (Masse et al., 2014). To measure **single unit activity (SUA**), time-varying voltages were manually ‘spike sorted’ by an experienced neurophysiologist using Plexon Offline Spike Sorter v3. This process identified action potentials belonging to putative individual neurons amongst the high amplitude voltage deviation events. Occasionally the same action potential can be recorded on multiple electrodes (this could happen if a neuron is very large, if an axon passes multiple electrodes, or if there is some electrical cross-talk in the recording hardware). To prevent creating duplicate single neuron units, we excluded ‘cross-talk units’ if their spike time series (using 1 ms binning) had greater than 0.5 correlation with another unit’s. When this happened, we kept the unit with the better spike sorting isolation. Unless otherwise stated, time-varying firing rate plots, also known as peristimulus time histograms (such as in **Figure 1B**) were constructed by smoothing spike trains with a 25 ms s.d. Gaussian kernel and averaging continuous-valued firing rates across trials of the same behavioral condition.

Spike sorting allows us to make statements about the properties of individual motor cortical neurons (for example, how many syllables they respond to, as in **Figure S3B**.) However, a limitation of spike sorting is that action potential event ‘clusters’ with insufficient isolation from other clusters are discarded. For chronic multielectrode array recordings, this can mean that activity recorded from the majority of electrodes is not analyzed, despite these neural signals having a strong relationship with the behavior of interest. This problem is particularly acute in human neuroscience, where replacing arrays, or using newer methods that provide a higher SUA yield (for example high density probes or optical imaging), is not currently possible. Analyzing voltage **threshold crossings (TCs),** i.e., relaxing the constraint that action potential events must be unambiguously from the same neuron, is an effective way to substantially increase the informational yield of chronic electrode arrays. In this study we examined TCs in a number of analyses. Decoding TCs or other non-SUA signals has become standard practice in the intracortical BCI field (e.g., (Ajiboye et al., 2017; Brandman et al., 2018; Collinger et al., 2013; Even-Chen et al., 2018; Pandarinath et al., 2017)). This method also provides information about the dynamics of the neural state (i.e., can be used to make scientific statements about ensemble activity under many conditions) despite combining spikes that may arise from one or more neurons; we provide empirical and theoretical justifications in (Trautmann et al., 2017). In the present study, when we refer to an ‘electrode’s’ firing rate, we mean TCs recorded from that electrode. When we refer to a neuron’s firing rate, we mean sorted single unit activity.

A threshold of −4.5 × root mean square (RMS) voltage was used for all analyses and visualizations except for the t-SNE visualization and decoding analyses shown in **Figure 3**. This threshold choice is somewhat arbitrary but is conservative; it accepts large voltage deviations indicative of action potentials from one or a few neurons near the electrode tip. For the **Figure 3** analyses, we used a more relaxed threshold of −3.5 × RMS because we found that this led to slightly better classification performance in a separate pilot dataset (consisting of T5 speaking five words and syllables, collected a month prior to the datasets reported here) which we used for choosing hyperparameters. The better performance of a less restrictive voltage threshold is consistent with collecting more information by accepting spikes from a potentially larger pool of neurons. This trade-off was acceptable because for these engineering-minded decoding analyses, we were less concerned about the possibility of missing tuning selectivity or fast firing rate details due to combining spikes from more neurons.

Electrodes with TC firing rates of less than 1 Hz (at a −4.5 × RMS threshold) were considered non-functioning and were excluded from analyses unless there was well-isolated SUA on the electrode. This electrode exclusion applied to both spikes and the local field potential signal described below. Electrodes having TCs time series with greater than 0.5 correlation with another electrode(s)’ were marked for cross-talk de-duplication. To determine which electrode to keep, we chose the one that had the fewest spikes co-occurring (1 ms bins) with the other electrode(s)’ (i.e., we kept the electrode with putatively more unique information).

For the neural decoding analyses (**Figure 3**) we also extracted a **high-frequency local field potential (HLFP)** feature from each electrode by taking the power of the voltage after filtering from 125 to 5,000 Hz (3^rd^ order bandpass Butterworth causal filtering forward in time). HLFP is believed to contain substantial power from action potentials (Waldert et al., 2013); we view this feature as capturing spiking “hash”, i.e., multiunit activity local to the electrode with contributions from smaller-amplitude and more distant action potentials than TCs. Our previous study found that this signal is highly informative about hand movement intentions and is useful for real-time BCI applications (Pandarinath et al., 2017). This feature has some similarities to the ‘high gamma’ activity examined by ECoG studies; the definition of high gamma varies in exact frequency from study to study, but generally has a lower cutoff between 65 and 85 Hz and an upper cutoff between 125 and 250 Hz (Bouchard et al., 2013; Chartier et al., 2018; Cheung et al., 2016; Dichter et al., 2018; Martin et al., 2014; Mugler et al., 2014; Ramsey et al., 2018). However, the intracortical HLFP in this study should not be viewed as being the exact same as ECoG high gamma activity due to differences in electrode location, electrode geometry, and HLFP’s higher frequency range.

### Task-related neural modulation

To quantify which electrodes’ spiking activity changed during speaking (**Figure 1A** insets, **Figure S3**), we calculated each electrode’s mean firing rate from 0.5 seconds before to 0.5 seconds after AO, yielding one datum per electrode, per trial. For each syllable, a rank-sum test was then used to determine whether there was a significant change in the distribution of single trial firing rates when speaking the syllable compared to the silent condition (p < 0.05, Bonferroni corrected for the number of syllables). To identify which electrodes responded to orofacial movements (**Figures 2, S4**) we performed a similar analysis, except that the analysis epoch was from 0.5 s before to 0.5 s after the go cue. This epoch captures strong modulation, as can be seen by the example firing rate plots in **Figure 2**. We note that firing rate changes preceding the go cue indicate either substantial movement preparation activity, or that the participants were “jumping the gun” and started moving in anticipation of the go cue; either way, this response indicates modulation related to making orofacial movements. In lieu of a silent condition, the movement conditions’ firing rate distributions were compared to that of the “stay still” condition. The same methods were used to quantify which single neurons’ spiking activity changed during speaking or orofacial movements; for this, we analyzed SUA rather than electrodes’ −4.5 × RMS TCs.

To summarize the differences in neural response magnitude following the audio prompt and following the go cue (**Figure S2C**), we first calculated the population response magnitude, which we defined as the mean across electrodes absolute value firing rate change compared to a baseline period from 1.25 s to 0.75 s before the trial start beep. We then calculated, for each speaking condition, the maximum value of this population firing rate change (i.e., the peak response magnitude across time) in a ‘prompt epoch’ (0.5 seconds before the prompt to 1 second after the prompt) and a ‘go epoch’ (0.5 seconds before the go cue to 2 seconds after the go cue). Firing rate change magnitude is always, by definition, a positive value, and therefore even if there was no speaking-related modulation, we would expect this metric to yield some non-zero value. To normalize for this floor effect, we subtracted the peak response magnitude for the silent condition (calculated in the same way as for the speaking conditions) from each syllable/word’s peak response magnitude. We then calculated the mean peak response across syllables/words. Lastly, we calculated the ratio of this mean peak response during the go epoch and during the prompt epoch.

### Breath-related neural modulation

To generate breath-triggered firing rates (**Figure S5**), we first identified breath peak times from the breath belt stretch transducer measurements. The belt signals were pre-processed by removing rare outlier values (> 50 μV difference between consecutive samples) and then low-pass filtering (3 Hz pass-band) the signal both forwards and backwards in time to avoid introducing a phase shift. An example of this filtered signal is shown in **Figure S5A**. Breath peaks were then found using the MATLAB *findpeaks* function, with key parameters of MinPeakDistance = 1 s, and MinPeakProminence = 0.3*B, where B is the median of all peak prominences found by first running *findpeaks* using MinPeakDistance = 5s (in other words, we required a peak to be at least 30% of the prominence of the “big” peaks in the data).

Breath peak-aligned firing rates were calculated by treating each identified breath peak as one trial, and trial averaging across neural snippets aligned to each breath peak time. Each TCs’ or SUA’s breath-related modulation depth was defined as the maximum – minimum firing rate observed in the interval from 2 seconds before the breath peak to 1.5 seconds after the breath peak. To calculate whether a given modulation depth was statistically significant, we used a shuffle control in which we compared the true data’s modulation depth to the distribution of modulation depths observed over 1,001 random shuffles in which faux peak breath times were uniformly drawn from the data duration. For comparing breath-related and speaking-related modulation depths (**Figure S5F**), we defined a given electrode’s speech modulation depth in the t5-syllables dataset as its maximum – minimum firing rate from 2.5 seconds before acoustic onset to 1 second after acoustic onset.

### Arm and hand versus orofacial and speaking movements comparisons

The neural ensemble modulation comparisons presented in **Figure S6** were calculated as follows: mean TC firing rates for each t5-comparisons dataset instructed movement condition were calculated for each electrode from 200 to 600 ms after the go cue. To account for baseline activity, the mean firing rates for the “do nothing” condition were subtracted from each movement condition’s activities. This resulted in a firing rate vector in high-dimensional (electrode) space. **Figure S6** reports this vector length for each movement condition.

### Single trial low-dimensional neural state projections

To visualize single-trial high dimensional neural data (**Figure 3A**) we used t-distributed stochastic neighbor embedding (tSNE), a dimensionally reduction technique which seeks to represent high-dimensional vectors (such as our time-varying, multielectrode neural data) in a low-dimensional space (such as a 2D plot that can be easily visualized). The tSNE algorithm finds a nonlinear mapping such that similar high-dimensional feature vectors end up close together in the low-dimensional view, while dissimilar vectors end up far apart (Van Der Maaten and Hinton, 2008). A neural feature vector was constructed for each trial as follows: for each functioning electrode, spike rates and HLFP power were calculated in ten 100 ms bins that spanned from 0.5 s before to 0.5 s after AO. These features were concatenated into a vector; for example, for the T5-syllables dataset, a single trial’s neural data were represented as a 104 electrodes × 2 features per electrode × 10 time bins = 2080-dimensional vector. All trials’ feature vectors were then projected into a 2D space using the *tsne* function in MATLAB R2017b’s Statistics and Machine Learning Toolbox with NumDimensions = 2; Perplexity = 15 (this is the number of local neighbors examined for each datum); Algorithm = exact (suitable for our relatively small dataset); and Standardize = true (this z-scores the input data, which was desirable due to the variability between different electrodes and the vastly different scales between spike rates and HLFP power). All other algorithm parameters were set to their defaults. **Figure 3A** does not have axis labels because t-SNE does not return meaningful axes or units; only the relative distances between points have meaning.

### Speech decoding

We evaluated how well the identity of the syllable or word being spoken could be decoded from neural data by classifying single trial neural data. Neural feature vectors were constructed for each trial as described above. These vectors were then associated with a class label, which was the sound being spoken (i.e., word, syllable, or silence). We trained support vector machines (SVMs), a standard classification tool, to predict the class label from a “new” neural feature vector which the classifier had not been trained on. Prediction accuracies were cross-validated using a leave-one-trial-out paradigm in which the classifier was trained on all trials except the trial being classified, and this was repeated for all trials in a dataset. Multiclass classification was achieved using the error-correcting output code (ECOC) technique, which trains multiple binary SVMs between all pairs of labels, i.e., a one-versus-one coding design. When classifying new input data, the ECOC technique picks the class that minimizes the sum of losses over the set of binary SVM classifiers. Specifically, we used MATLAB R2015a’s implementation: a multiclass model object was fit (*fitcecoc*) using the SVM template (*templateSVM*). Key parameters were to use a linear kernel; OutlierFraction = 0.05 (expecting 5% of data points to be outliers); and Standardize = true (which z-scores the neural features based on the training data). All other parameters were set to their default values. We note that we did not heavily optimize our classification method; rather, our goal here was to use a standard tool to gauge the classification performance that these intracortical neural signals support. More sophisticated techniques from machine learning (e.g., (Angrick et al., 2018; Livezey et al., 2018)) are likely to provide additional improvements.

To measure chance prediction performance, we used a shuffle test in which we randomly permuted the class labels associated with all trials’ neural data. The same classifier training and leave-one-out prediction process was then repeated on these shuffled data 101 times.

### Neural population dynamics

An underlying motivation for the neural population dynamics analyses described in the next several sections is the idea that the activity of many thousands or millions of neurons in a circuit (of which we can only measure on the order of 100 in humans with current technology) can be summarized by the time-varying activity of a handful of latent ‘factors’. In this framing, individual neurons’ firing rates reflect various mixtures of these underlying factors; in all of the analyses we used, this mapping from factors to firing rates is assumed to be linear. These factors are not meant as discrete physical “things” in the brain, but rather are mathematical abstractions which capture meaningful patterns in the behavior of networks of neurons. They are useful insofar as they can generate hypotheses about the computations being performed. To this end, not only can latent factors succinctly describe the ‘neural state’ (i.e., the firing rate of all neurons at a given moment in time), but furthermore, the time evolution of these factors is often more conducive to interpretation and understanding than more complex descriptions of all the individual neurons’ firing rates.

Here, for example, we build on previous studies showing that these factors’ changes over time can be effectively modeled as a lawful time-varying oscillatory dynamical system (Churchland et al., 2012), and that they reveal a simple population-level pattern in which there is a stereotyped response at the initiation of many different movements (Kaufman et al., 2016). This ‘dynamical system’ framework is extensively reviewed in (Shenoy et al., 2013) as well in the two key studies that inspire the neural population dynamics analyses of the present study (Churchland et al., 2012; Kaufman et al., 2016). We looked for the aforementioned dynamical motifs using two different dimensionality reduction techniques that were specifically designed to reveal the presence (or absence) of these population dynamics features.

For these analyses, we examined the prompted word speaking task datasets because this was a more naturalistic behavior than the prompted syllable speaking task. Participants reported that it was more difficult to discriminate syllables than words, and that speaking stand-alone syllables felt somewhat awkward; they expressed doubt about whether they were saying the syllables correctly, whereas saying words was easy. Consequently, a practical benefit of the words task over the syllables task is that behavior was more stereotyped across trials, which facilitates trial-averaging, and there were very few mis-heard or mis-spoken words. Unlike for the syllables task, in the words task both participants had close to 30 trials each for all ten speaking conditions.

Both of these sets of neural population state analyses were performed on TCs, which contained more information about the neural population state than the more limited number of recorded SUA. All electrodes with TC firing rates greater than 1 Hz were included. The Churchland-Cunningham and Kaufman studies analyzed a combination of both SUA from single-electrode recordings and TCs from multielectrode recordings, depending on the dataset, while (Pandarinath et al., 2015) also analyzed just TCs. To avoid cumbersome switching of terms when describing our methods and comparing them to those of these previous studies, we will use the generic term ‘unit’ to refer to a single channel of neural information, whether it be SUA or TCs.

### Condition-invariant signal

The first population dynamics motif we tested for was a specific form of population-level structure at the initiation of movement: a large condition-invariant signal, previously described by Kaufman and colleagues (Kaufman et al., 2016). We closely followed Kaufman’s analysis methods, adapting them as necessary for these human speaking datasets. As in (Kaufman et al., 2016), spike trains were trial-averaged within a behavioral condition (in our case, speaking one of the ten different words), smoothed with a 28 ms s.d. Gaussian, and ‘soft normalized’ with a 5 Hz offset. Normalization means that each unit’s firing rate was normalized by its range across all times and conditions. This prevents units with very high firing rates from dominating the estimate of neural population state. The ‘soft’ refers to adding an offset (5 Hz in these analyses) to the denominator to reduce the influence of units with very small modulation. Trial-averaged firing rates were calculated from 200 ms before go cue to 400 ms after the go cue in order to focus on the epoch when speech production was being initiated. This yields a N × C × T data tensor, where N is the number of units, C is the number of word conditions (10), and T is the number of time samples (600, using 1 ms sliding bins).

We used demixed principal components analysis (dPCA), a dimensionality-reduction technique developed by Kobak, Brendel and colleagues (Kobak et al., 2016), to look for condition-invariant activity patterns in these high-dimensional neural recordings. This dimensionality reduction method is conceptually similar to PCA, in that it finds a specified number of dPC ‘components’ that can be thought of as “building blocks” from which the responses of individual units can be composed. As with PCA, dPCA attempts to compress the data by identifying dimensions that capture a large fraction of the variance. This takes advantage of the fact that unless the responses of neurons are all independent from one another (which in practice is not the case), then most of the variance of the full population response can be accurately reconstructed as a weighted sum of a smaller number of dPC components. Where dPCA differs from PCA is that it can explicitly attempt to find components that marginalize variance attributable to different parameters of the experiment (such as time or task variables). This is possible because dPCA is a supervised method that trades off finding dimensions that maximize variance in favor of finding dimensions that partition the variance based on labeled properties of the data.

In our case, this ‘demixing’ was attempted between: 1) condition and condition + time interactions, which together form the condition-dependent (CD) components of the neural population activity; and 2) time only, which form condition-invariant (CI) components. In other words, dPCA sought a set of components of the population activity for which the time-varying neural responses during producing different words look the same, and also for another set of components which vary across speaking conditions (i.e., are “tuned” for what word is being spoken). Importantly, such variance marginalization (i.e., demixing the parameters) may not be achievable; it depends on the structure of the data itself. Each component that dPCA returns is associated both with how much overall neural variance it captures (the lengths of the bars in **Figure 4A**), and how much of this variance is CI or CD (red and blue fraction of each bar, respectively). Thus, the success of this demixing can be examined based on how purely CI or CD each component is. This in turn reveals whether there exists a large and almost completely condition-invariant component of the population neural activity.

Kaufman and colleagues used an earlier version of the dPCA method and code package, called ‘dPCA-2011’ (Brendel et al., 2011). We used the MATLAB implementation of ‘dPCA-2015’ (Kobak et al., 2016), downloaded from http://github.com/machenslab/dPCA. This is an updated, improved, and widely adopted version of the technique which was not yet available at the time when the (Kaufman et al., 2016) analyses were performed. We specified that dPCA should return eight total components, which was less than then 10 to 12 used in (Kaufman et al., 2016). This reflects the reduced complexity of our datasets, in the sense that they had fewer conditions (10 versus 27-108) and fewer units (96-106 versus 116-213). We also repeated the analyses using 5 to 12 dPCs and observed very similar results. Default *dpca* function parameters were used.

Unlike the dPCA-2011 used by (Kaufman et al., 2016), dPCA-2015 does not enforce that the neural dimensions found for capturing variance attributable to different parameters (here, the CI and CD components) be orthogonal. For example, while the three different CI components for T5 in **Figure 4A** are orthogonal by construction (as are the five different CD components), the CI and CD components need not be orthogonal. We therefore quantified the degree of orthogonality between the CIS_1_ component and the CD components by measuring the principal angle between CIS_1_ and the subspace defined by CD components. Specifically, we used the *subspacea* package for MATLAB, downloaded from https://www.mathworks.com/matlabcentral/fileexchange/55-subspacea-m (Knyazev and Argentati, 2002).

### Rotatory dynamics

The second form of neural population structure we tested for was rotatory (i.e., oscillatory) low-dimensional dynamics. We applied methods previously developed to identify and quantify rotatory dynamics in motor cortex during NHP arm reaching (Churchland et al., 2012). These methods were also recently applied to show rotatory dynamics during hand movements of BrainGate2 study participants (Pandarinath et al., 2015). Churchland, Cunningham and colleagues introduced the jPCA dimensionality reduction technique for this purpose; we employed their MATLAB analysis package, downloaded from https://churchland.zuckermaninstitute.columbia.edu/content/code.

Trial-averaged firing rates for each word speaking condition were generated from 150 ms before to 100 ms after acoustic onset to capture an epoch when speech-producing articulator movements were being produced. Following (Churchland et al., 2012; Pandarinath et al., 2015), these firing rates were soft-normalized with a 10 Hz offset and smoothed with a Gaussian kernel; we used a 30 ms s.d. kernel as in (Pandarinath et al., 2015). These firing rates were ‘centered’ by subtracting the across-condition mean firing rate of each unit at each time point, and then sampled every 10 ms. The dimensionality of these data was reduced via PCA to six; this ensured that rotatory dynamics would be sought within population activity components that were strongly present in the data. jPCA was then used to find planes with rotatory structure within this six-dimensional subspace. The jPCs are found by fitting the following linear dynamical system:

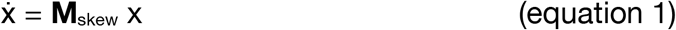

where x is the neural state (i.e., the PCA dimensionality-reduced population firing rate) at a given time, 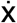 is its time derivative, and **M**_skew_ is constrained to be a skew-symmetric matrix. The first jPC plane, which has the strongest rotatory dynamics, is defined by the two complex eigenvectors of **M**_skew_ with the largest eigenvalues. The choice of real vectors jPC_1_ and jPC_2_ within this plane is arbitrary and, following convention, were chosen such that conditions’ activities are maximally spread along jPC_1_ at the start of the analysis epoch. **Figures 5A** and **S5** plot the trial-averaged population activity during speaking each word (after subtracting the across-conditions mean) in this top jPC plane. The red/black/green color of each word condition’s neural trajectory corresponds to its projection along jPC_1_ at the start of the epoch; this display style is intended to assist in observing that amplitude and phase tend to unfold lawfully from the initial neural state. It is worth emphasizing that each jPC is simply a linear weighting of different units’ firing rates, and that the six jPCs form an orthonormal basis set that spans the same subspace as the top six PCs. The strength of rotatory dynamics was quantified as the goodness of fit for equation 1 for a 2×2 **M**_skew_ in the first jPCA plane, and for a 6×6 **M**_skew_ in the 6-dimensional subspace defined by the top 6 PCs of the data. **Figure 5B** reports this 6D fit quality.

### Statistical testing of rotatory dynamics

To calculate the statistical significance of rotatory population dynamics structure in our data, we applied the ‘neural population control’ approach developed by Elsayed and Cunningham (Elsayed and Cunningham, 2017). This method was developed to address a potential concern that many specific phenomena that an experimenter could test for (such as fitting low-dimensional rotatory dynamics to neural data) can be found “by chance” in a sufficiently high-dimensional, complex dataset such as the time-varying firing rates of many neurons. To address this, the method tests whether an observed feature of the population activity is “novel” in the sense that it cannot be trivially predicted from known simpler features in the data. This is achieved by constructing surrogate datasets with simple population structure (in the form of means and correlations across time, neurons, and behavioral conditions) matched to the real data. If the neural recordings contain population-level structure that is coordinated above and beyond these first and second-order features, then the quantification method used to describe this structure should return a stronger read-out when applied to the original dataset than to the surrogate datasets.

In our case, we used this approach to test whether it is “surprising” to see rotatory dynamics in neural population data, given the particular smoothness across time, units, and word speaking conditions present in these data. A similar approach was used in (Elsayed and Cunningham, 2017) to further validate the original rotatory dynamics finding of (Churchland et al., 2012). We used the MATLAB code associated with (Elsayed and Cunningham, 2017) from https://github.com/gamaleldin/TME to generate 1,000 surrogate datasets with time, neuron, and condition means and covariance matched to the real data using the tensor maximum entropy algorithm (‘surrogate-TNC’ flag in *fitMaxEntropy*). We then ran the same jPCA analyses described above on these surrogate datasets and recorded the rotation dynamics goodness of fit for the best **M**_skew_ matrix found for each surrogate dataset. This distribution of surrogate dataset R^2^ values serves as a null distribution for significance testing: we calculated a *P* value by counting how many of the surrogate datasets’ R^2^ exceeded that of the true original dataset.

## SUPPLEMENTAL MATERIALS

**Figure S1.**
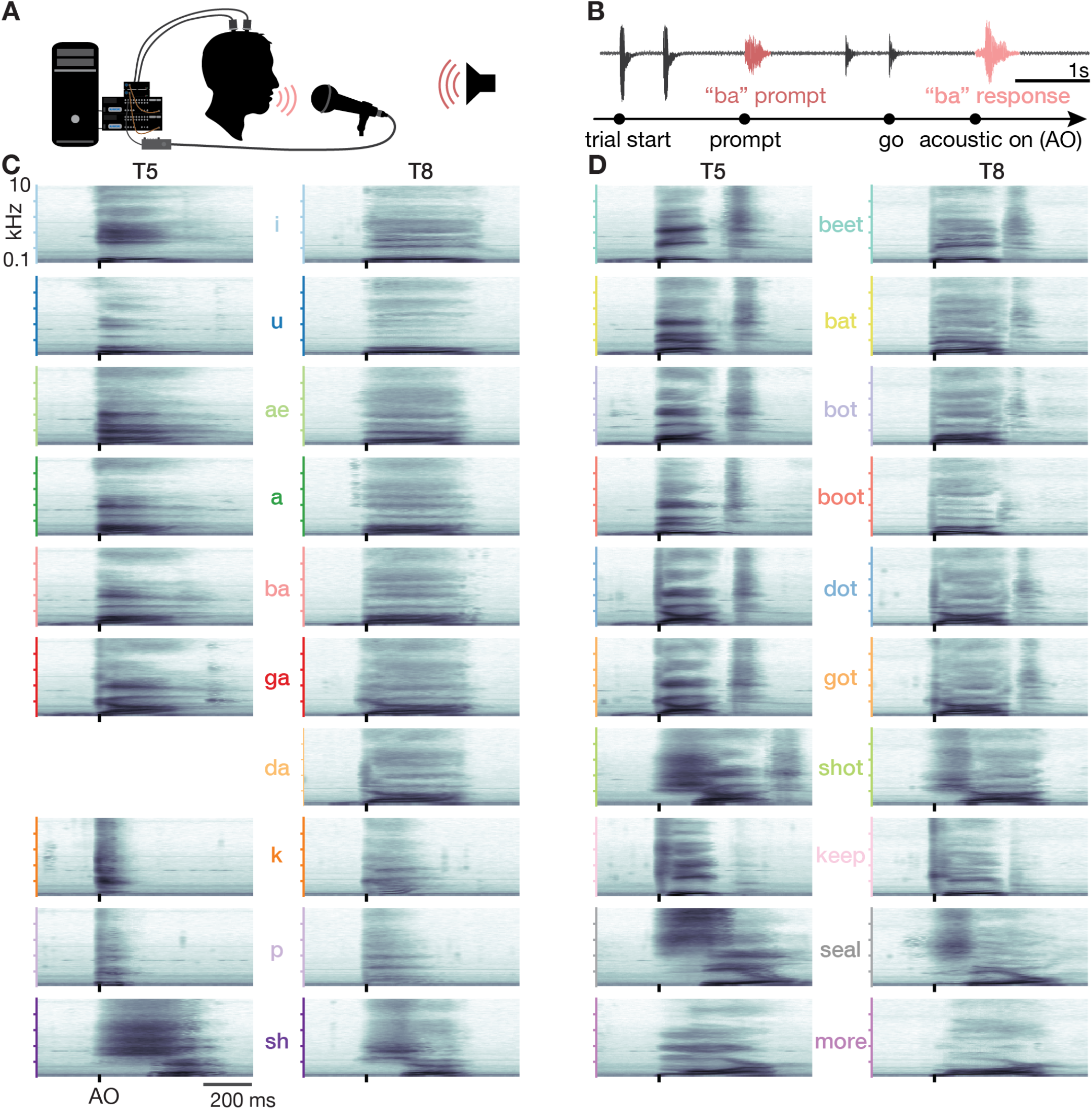
Prompted speaking task. (A) Schematic of the experiment setup. Participants performed a prompted speaking task in which they heard a syllable or word played from a computer speaker. They were instructed to speak back that sound after hearing a go cue. Motor cortical neural signals were recorded during the task. A microphone captured both the prompts and participant’s speech. The microphone recording was amplified and captured as an analog input by the neural signal processor, thus synchronizing the audio data with the neural data. (B) Example acoustic waveform recorded during one trial (top) and the trial’s corresponding task event timeline (bottom). The syllable prompt played by the computer speaker and the subsequent response spoken by the participant are colored in pink. Two beeps indicated the start of a trial, and the second of two clicks was the go cue that instructed the participant to repeat back the prompted sound. AO is acoustic onset. (C) Acoustic spectrograms for the participants’ spoken syllables. Power was averaged over all analyzed trials. Note that *da* is missing for T5 because he usually misheard this cue as *ga* or *ba*. (D) Same as panel c but for the T5-words and T8-words datasets.

**Figure S2.**
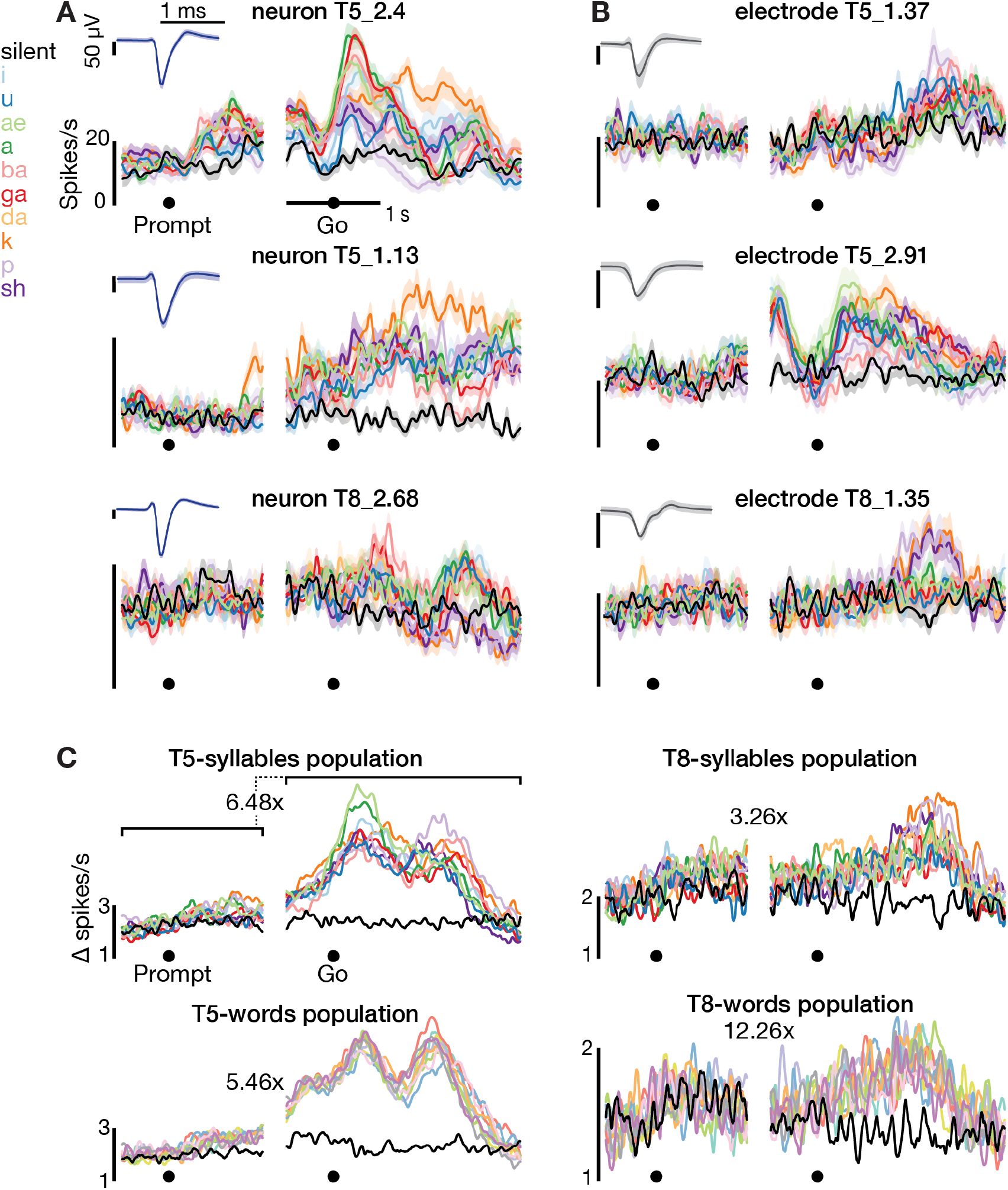
Motor cortical modulation is greater during speaking than hearing speech prompts. (A) Firing rates for the same neurons shown in **Figure 1B** are shown here aligned to the auditory prompt (when the syllable was played to the participant via computer speaker) as well as to the go cue that instructed the participant to speak. (B) Many electrodes recorded action potential events that could not necessarily be attributed to an individual neuron based on their waveforms (i.e., could not be spike-sorted). Nonetheless, these threshold-crossing spikes (TCs) exhibited speech-related activity. We therefore included them in our analyses to maximize the available information about the motor cortical population ensemble. Firing rates for three example electrodes’ TCs (at a voltage threshold of −4.5 x RMS) are shown. Insets show these TCs’ spike-triggered waveforms in the same format as the sorted single neurons’ waveforms. (C) Population activity change from baseline for all four speech datasets. Audio Prompt-and Go-aligned firing rates were calculated for all functioning electrodes’ −4.5 × RMS TCs. We then subtracted each electrodes’ ‘baseline activity’ from these responses to yield a time-varying firing rate change. Each trace shows, for a given syllable or word, the mean absolute value firing rate change across electrodes. Colors corresponding to each word are the same as in **Figure S1**. Baseline was defined as the firing rate during the inter-trial period (specifically, from 1.25 to 0.75 seconds before the beep indicating the start of a trial). Above each plot we report the ratio of maximum modulation (mean across sounds) during the go versus prompt epochs.

**Figure S3.**
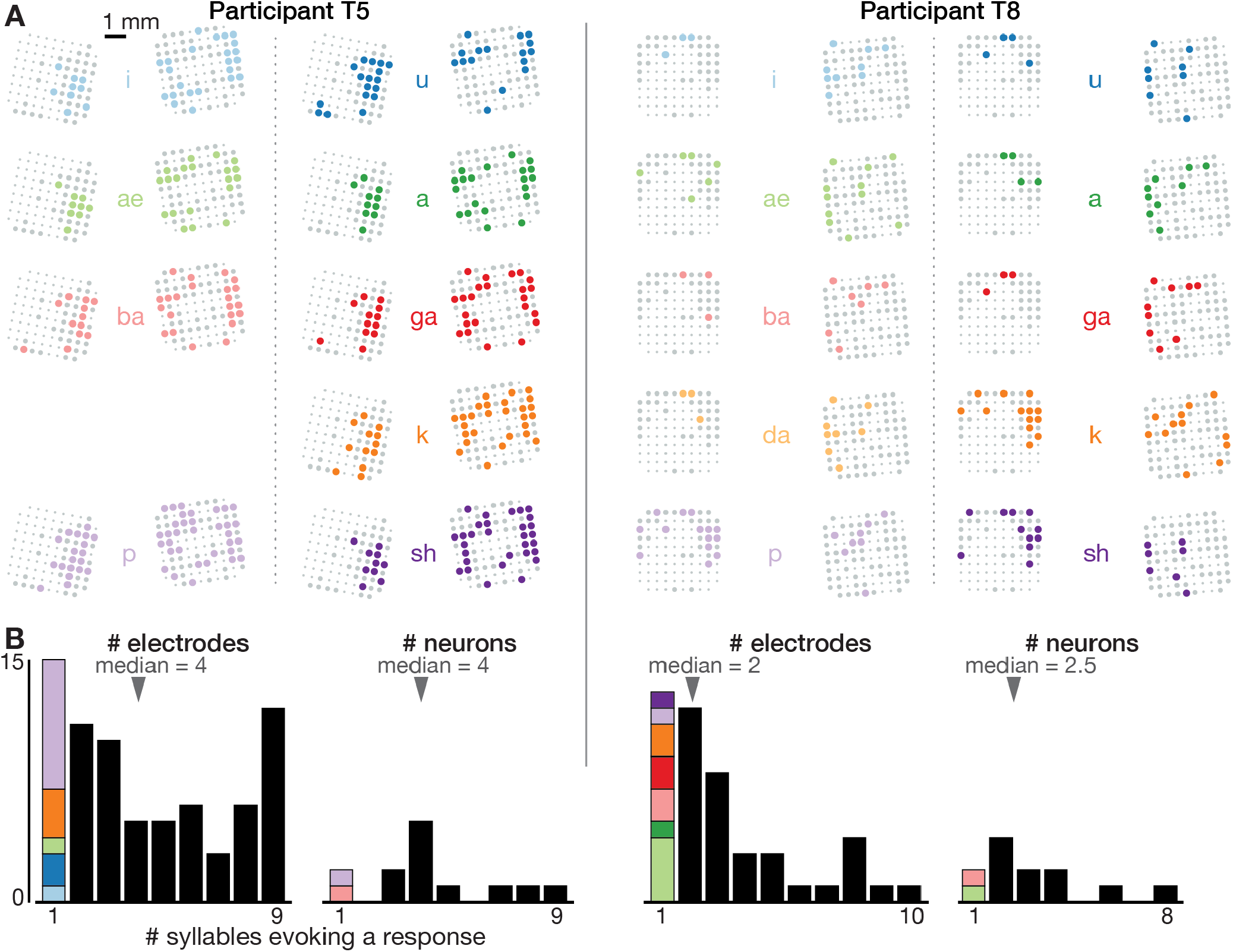
Neural correlates of spoken syllables are not spatially segregated in dorsal motor cortex. (A) Electrode array maps similar to **Figure 1A** insets are shown for each syllable separately to reveal where modulation was observed during production of that sound. Electrodes where the TCs firing rate changed significantly during speech, as compared to the silent condition, are shown as colored circles. Non-responding electrodes are shown as larger gray circles, and non-functioning electrodes are shown as smaller dots. Adding up how many different syllables each electrode’s activity modulates in response to yields the summary insets shown in **Figure 1A**. These plots reveal that electrodes were not segregated into distinct cortical areas based on what syllables they responded to. (B) Histograms showing the distribution of how many different syllables evoke a significant firing rate change for electrode TCs (each participant’s left plot) and sorted single neurons (right plot). The first bar in each plot, which corresponds to electrodes or neurons whose activity only changes when speaking one syllable, is further divided based on which syllable this response was specific to (same color scheme as in panel A). This reveals two things. First, single neurons or TCs (which may capture small numbers of nearby neurons) were typically not narrowly tuned to one sound. Second, there was not one specific syllable whose neural correlates were consistently observed on separate electrodes/neurons from the rest of the syllables.

**Figure S4.**
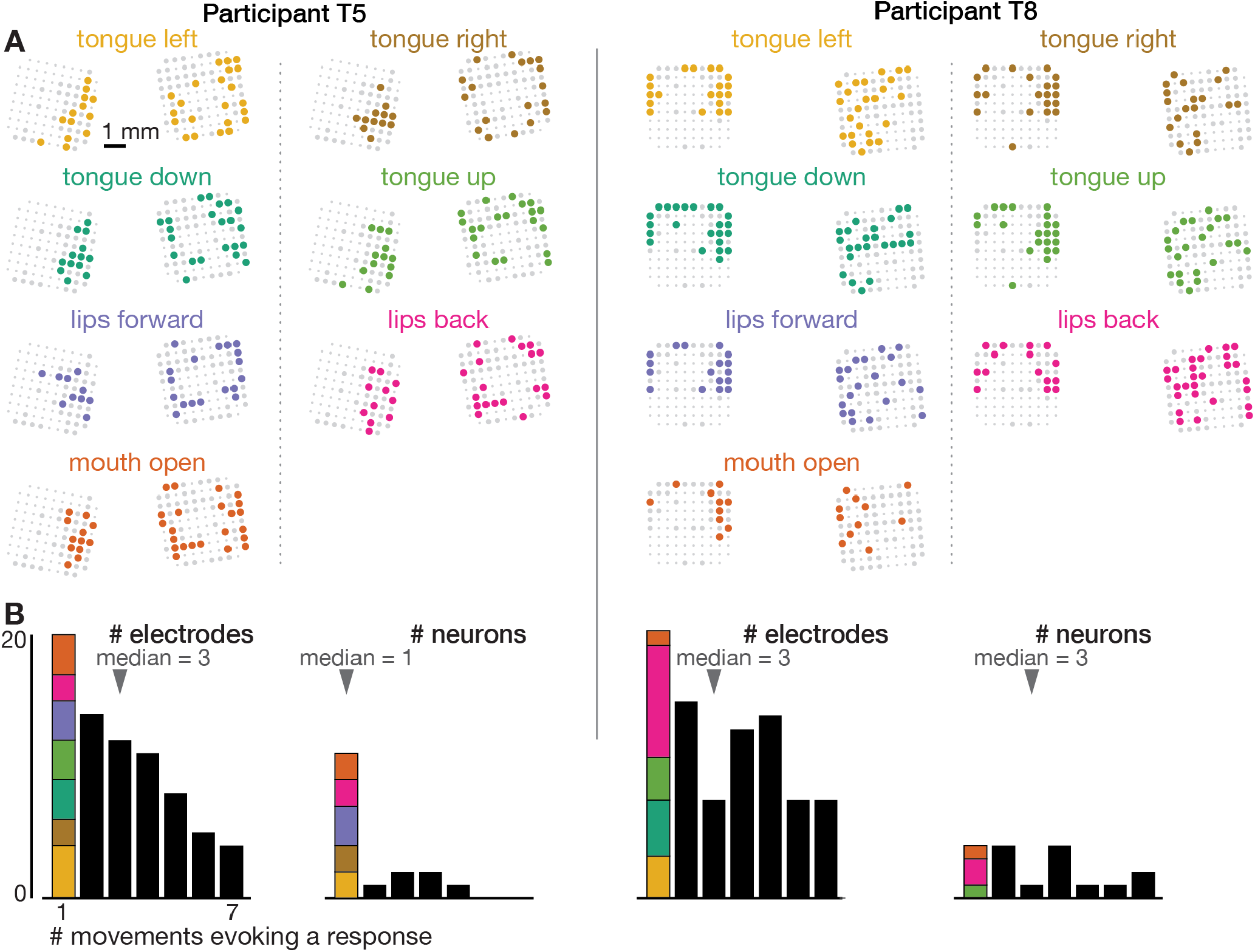
Neural correlates of orofacial movements are not spatially segregated in dorsal motor cortex. The orofacial movements data were analyzed and are presented analogously to the speaking data in **Figure S3**.

**Figure S5.**
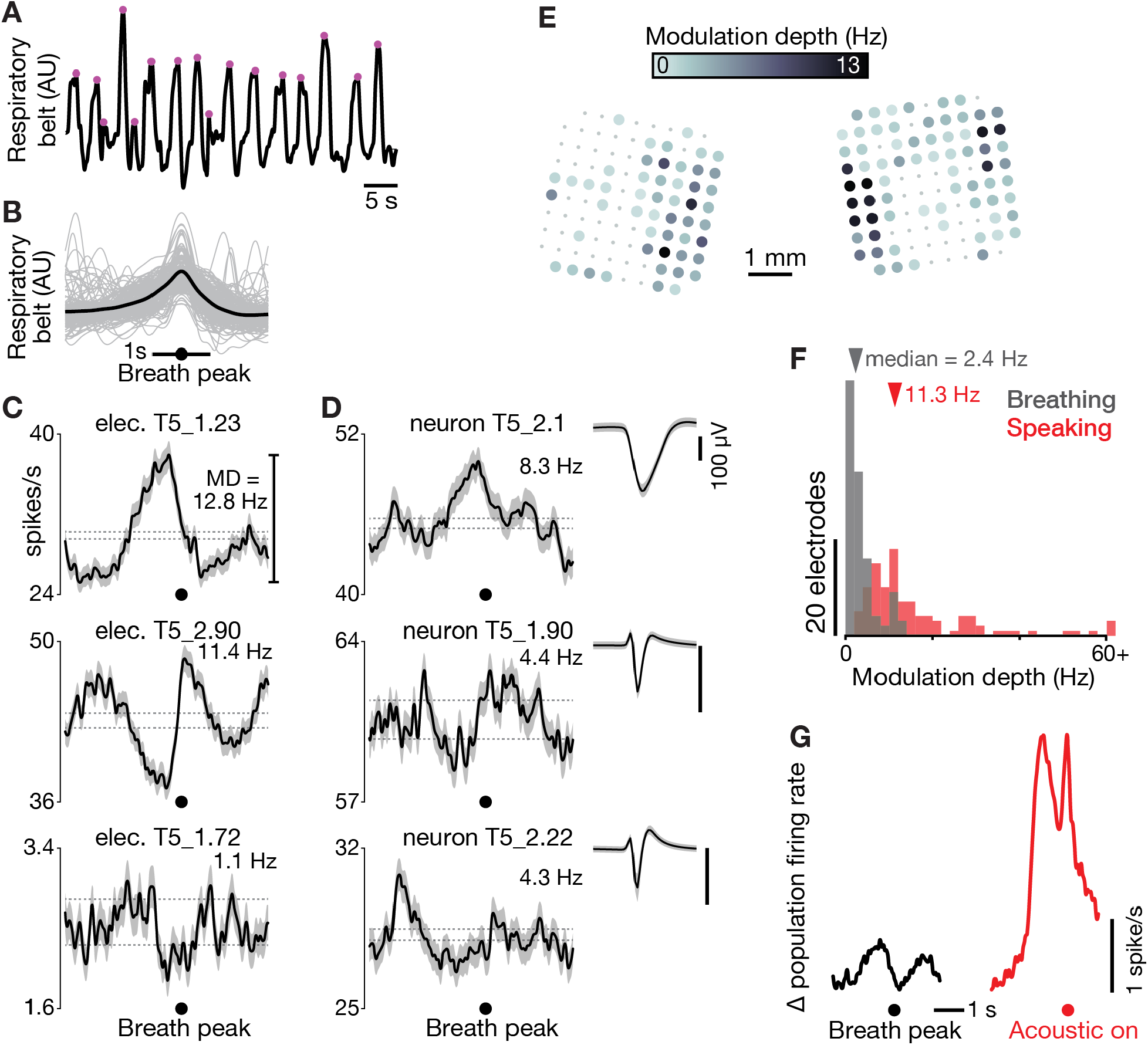
Dorsal motor cortex correlates of breathing. (A) We recorded both neural data and a breathing proxy (the stretch of a belt wrapped around participant T5’s abdomen) while he performed a BCI cursor task or sat idly. An example of continuous breath belt measurements is shown. Violet dots mark the identified breath peak times. (B) Gray traces show example respiratory belt measurements for 200 breaths, aligned to the breath peak. The mean of the 727 breaths in the t5-breathing dataset is shown in black. (C) Breath-aligned firing rates for three example electrodes’ TCs (mean ± s.e.m). Breath-related modulation depth was calculated as the peak-to-trough firing rate difference. Horizontal dashed lines show the p=0.001 modulation depths for a shuffle control in which breath peak times were chosen at random. (D) Breath-aligned firing rates were also calculated for sortable SUA, whose spike waveform snippets are shown as in **Figure 1B**. Three example neurons with large waveforms are shown, but all 18 sorted units had significant modulation. This argues against the breath-related modulation being an artifact of breath-related electrode array micromotion causing a change in TC firing rates by bringing additional units in or out of recording range. (E) Breath-related modulation depth for each functioning electrode’s TCs, presented as in **Figure 1A**. (F) Histograms of breath-related modulation depths (gray) and speaking-related modulation depths (red) for all functioning electrodes in the t5-breathing and t5-syllables datasets, respectively. Two outlier datums are grouped in a > 60 Hz bin. Dorsal motor cortical modulation was much stronger for speaking than breathing. (G) Mean population firing rates (averaged across all functioning electrodes) are shown during breathing (black), aligned to breath peak, and during speaking (red), aligned to acoustic onset. The vertical offsets of each trace have been bottom-aligned to facilitate comparison.

**Figure S6.**
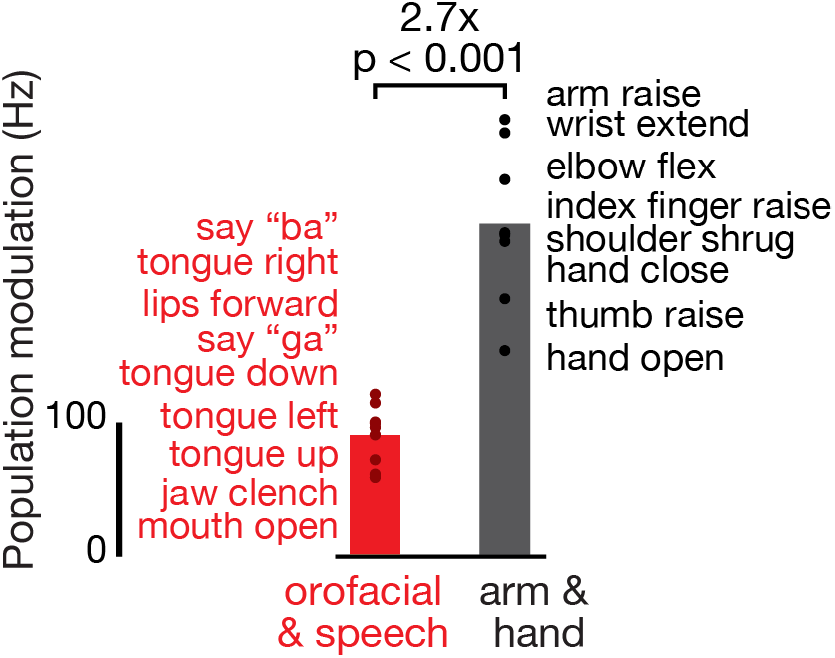
Dorsal motor cortex modulates more strongly during attempted arm and hand movements than orofacial movements and speaking. Participant T5 performed a variety of instructed movements. We compared the neural responses when attempting orofacial and speech production movements (red), versus when attempting arm and hand movements (black). Note that T5, who has tetraplegia, was able to actualize all of the orofacial and speech movements, but not all of the arm and hand movements. Each point corresponds to the mean TCs firing rate modulation for a single instructed movement (movements are listed in the order corresponding to their points’ vertical coordinates) in the t5-comparisons dataset. Modulation strength is expressed as a Euclidean distance in firing rate space (one dimension per functioning electrode) between the ensemble firing rates during attempting the instructed movement, and the ensemble firing rates during otherwise similar trials where the instruction was to “do nothing”. Bars show the mean modulation across all the movements in each grouping. A rank-sum test was used to compare the distributions of orofacial & speech and arm & hand modulations.

**Figure S7.**
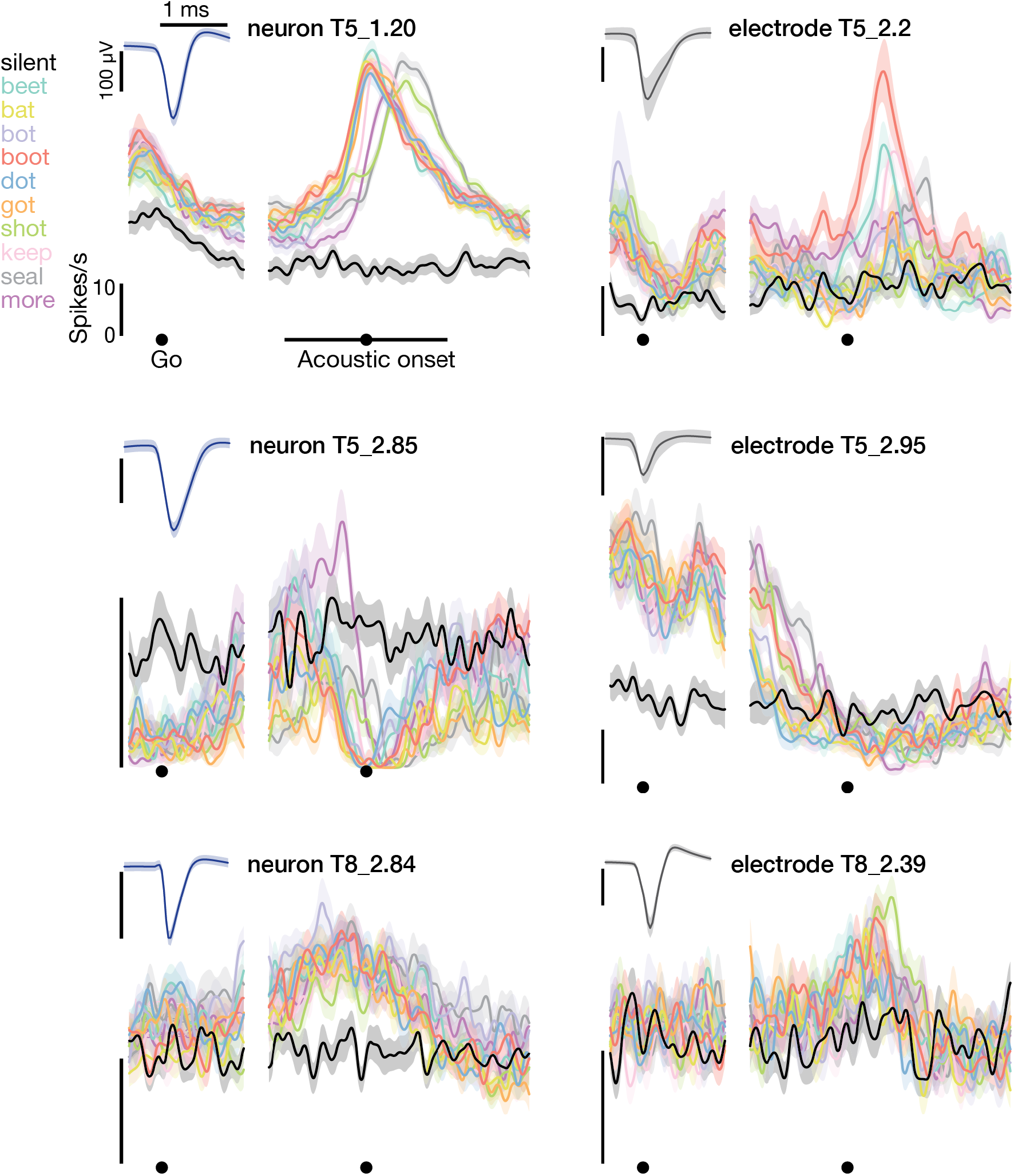
Example neural activity while speaking short words. Firing rates during speaking of short words for three example neurons (blue spike waveform insets) and three example electrodes’ −4.5 × RMS threshold crossing spikes (gray inset waveforms). Data are presented similarly to **Figure 1B**.

**Figure S8.**
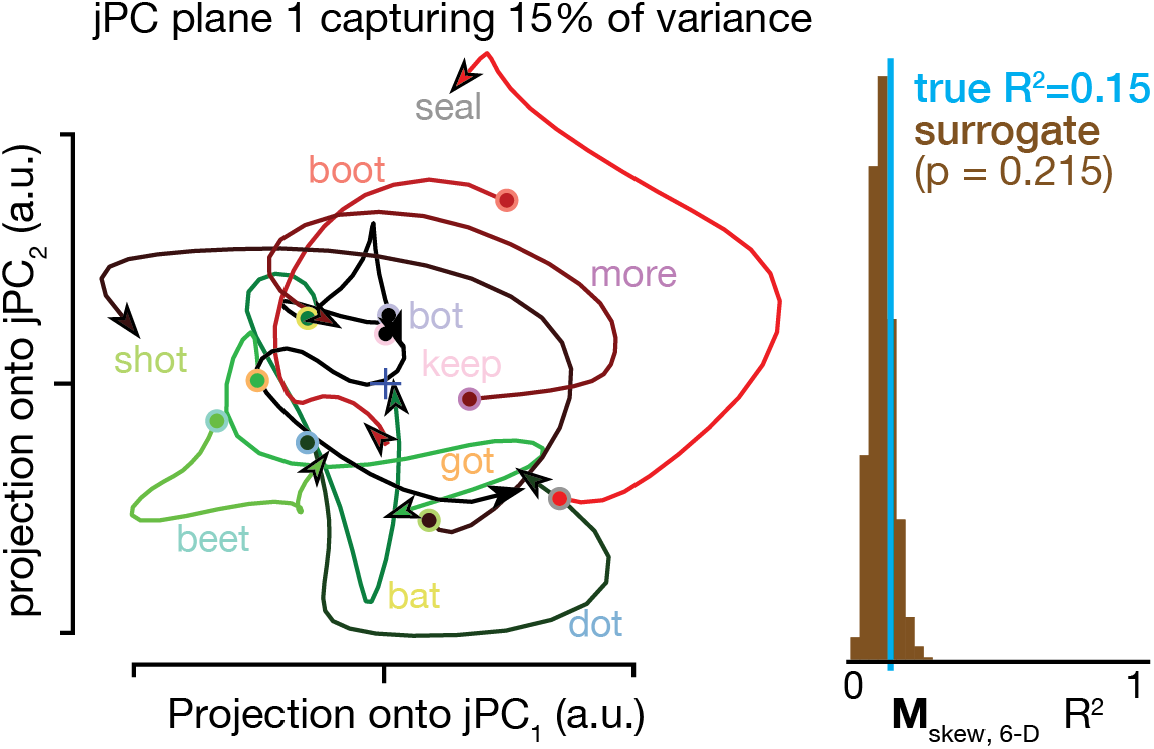
Rotatory dynamics for participant T8. Participant T8’s neural state trajectories projected into the first jPCA plane (left), and statistical testing of whether the goodness of fit of rotatory dynamics exceeds that of surrogate datasets (right). Data are presented as in **Figure 5**.

## Supplemental Movie 1

(duration: 1m 12s) Example audio and neural data from eleven contiguous trials of the prompted syllables speaking task. The audio track was recorded during the experiment and digitized alongside the neural data; it starts with the two beeps indicating trial start, after which the syllable prompt was played from computer speakers, followed by the go cue clicks, and finally the participant speaking the syllable. The video shows the concurrent −4.5 × RMS threshold-crossing spikes rate on each electrode. Each circle corresponds to one electrode, with their spatial layout corresponding to electrodes’ location in motor cortex as in the **Figure 1A** inset. Each electrodes’ color represents its firing rate (soft-normalized with a 10 Hz offset, smoothed with a 50 ms s.d. Gaussian kernel), with the color map going from pink (minimum rate) to yellow (maximum rate). Non-functioning electrodes are shown as small gray dots. Data from the T5-syllables dataset, trial set #23.

